# The tunicate metabolite 2-(3,5-diiodo-4-methoxyphenyl)ethan-1-amine targets ion channels of vertebrate sensory neurons

**DOI:** 10.1101/2021.05.05.442828

**Authors:** Noemi D. Paguigan, Yannan Yan, Manju Karthikeyan, Kevin Chase, Jackson Carter, Lee S. Leavitt, Albebson L. Lim, Zhenjian Lin, Tosifa Memon, Sean B. Christensen, Bo H. Bentzen, Nicole Schmitt, Christopher A. Reilly, Russell W. Teichert, Baldomero M. Olivera, Shrinivasan Raghuraman, Eric W. Schmidt

**Author notes:** Corresponding Author **Eric W. Schmidt** –. E- mail., **Shrinivasan Raghuraman** –.

## Abstract

Marine tunicates produce defensive amino-acid derived metabolites, including 2-(3,5-diiodo-4-methoxyphenyl)ethan-1-amine (DIMTA), but their mechanisms of action are rarely known. Using an assay-guided approach, we found that out of the many different sensory cells in the mouse dorsal root ganglion (DRG), DIMTA selectively affected low-threshold cold thermosensors. Whole-cell electrophysiology experiments using DRG cells, channels expressed in *Xenopus* oocytes and human cell lines revealed that DIMTA blocks several potassium channels, reducing the magnitude of the afterhyperpolarization and increasing the baseline [Ca2+]i of low-threshold cold thermosensors. When injected into mice, DIMTA increased the threshold of cold sensation by >3 oC. DIMTA may thus serve as a lead in the further design of compounds that inhibit problems in the cold-sensory system, such as cold allodynia and other neuropathic pain conditions.

## INTRODUCTION

The sensory nervous system consists of many cell classes of glia and neurons that collect and integrate information about the environment. In one of the major vertebrate sensory nerve centers, the dorsal root ganglion (DRG) contains many different types of peripheral nerve cells, which relay signals including temperature, position/balance, touch, itch, and pain.^1-3^ While the individual ion channels and receptors found in these cells are widely distributed in the body, the combination of those receptors and channels is unique to each cell type and leads to the ability to selectively sense different signals.^4^ The challenge in neuronal drug discovery could therefore be viewed as how to target specific cell types, rather than how to modulate individual proteins that are likely to be more widely distributed.

To meet this challenge, we applied constellation pharmacology, a calcium fluorescence imaging method of distinguishing cell types in different tissues on the basis of their response to pharmacological agents, and of discovering drugs that selectively target cell types.^5^ Here, we describe a compound from a marine tunicate that at lower doses selectively targets low-threshold cold thermosensors (LTCTs) from DRG neurons, likely by inhibiting a combination of potassium channels specifically expressed in these neurons. Cold sensation in the periphery is mediated by at least three cell types of DRG neurons: LTCTs, high-threshold cold thermosensors (HTCTs) and cold nociceptors (CNs). CNs are primarily responsible for causing pain in response to extreme and potentially damaging cold, while LTCTs provide fine information about temperature variations.^6^ Differences in the combination of ion channels present in cold sensory neurons define their roles, and genetic or pharmacological modulation of the ion channels can change the threshold of cold sensation in animals. Moreover, injuries to nerves can differentially change the gene-expression profiles in different cell types, resulting in the development of cold allodynia and other neuropathic pain disorders.^7, 8^ Thus, pharmacologically targeting selective cell types can be an alternative approach to drug discovery for pain and other neurological disorders.

Constellation pharmacology has been used to identify targets of bioactive peptides, but its application to small molecules has been significantly more challenging, in large part because of the relatively promiscuous nature of small molecule drugs.^9^ In our ongoing efforts to discover new therapeutics for the management of chronic pain, we screened a library of animal-derived marine natural products to identify small molecules that modulate signaling in the mouse DRG neurons. We screened the extracts of *Didemnum* sp. tunicates using constellation pharmacology of DRG neurons, leading to the discovery of a small molecule 2-(3,5-diiodo-4-methoxyphenyl)ethan-1-amine (diiodomethoxytyramine; DIMTA) as an ion channel blocker. DIMTA is one of the major metabolites of several tunicates, where it was reported to be modestly antifungal and cytotoxic.^10-13^ Its neuromodulatory and ion channel blocking properties were not known. Here, we showed that DIMTA activates LTCTs. Like most thermosensors, LTCTs are activated by the transient receptor potential M8 (TRPM8), also known as the menthol receptor. We screened DIMTA against TRPM8 and several potassium channels, showing that it does not modulate TRPM8 but instead inhibits potassium channels. In addition, constellation pharmacology data suggest that DIMTA is a relatively promiscuous voltage-gated ion channel (VGIC) blocker at high doses (25 µM), but at lower doses (2.5 µM) it shows selectivity for several potassium channels.

In mammalian tissues, there are about 70 distinct genes encoding for several types of K^+^ channels including voltage-gated (K_V_) and Ca^2+^-activated (K_Ca_) K^+^ channels, “leak” K^+^ channels and the inward rectifier (K_ir_) K^+^ channels.^14^ These channels perform a variety of physiological functions ranging from regulating the cell’s resting membrane potential and excitability to controlling neurotransmitter release. Additional complexity arises because many K-channels are heteromeric proteins. Because the total number of K-channel heteromeric combinations is unknown but may be substantial, it is impossible to exhaustively screen all possible K-channel targets. Here we describe a simplified approach to identify K-channel targets by focusing on the K-channel transcripts found in cell types that were selectively targeted by DIMTA. By comparing the transcripts (and electrophysiological analyses) with cell types that were unaffected, we narrowed down the list of K-channel targets. Modulating the functions of these channels represents an immense potential for the development of antinociceptive pharmacology for the management of chronic pain and other conditions.

While DIMTA was relatively promiscuous on potassium channels, it was potent and selective in targeting specific LTCT neurons. This cell-type selectivity (in contrast to molecular target selectivity) prompted us to investigate whether DIMTA would affect cold sensation in mice. We hypothesized that depolarization (activation) of LTCTs would increase cold sensitivity in mice, but strikingly DIMTA instead significantly inhibited the response to cold, decreasing the cold temperature sensitivity by more than 3°C. DIMTA thus provides a new lead scaffold for the design of cell-type selective potassium channel blockers, and potentially for treatments for cold allodynia resulting from clinical neuropathic pain states.

## RESULTS AND DISCUSSION

### DIMTA elevates intracellular calcium in LTCTs

Assay-guided screening using a panel of marine animal extracts led to the identification of DIMTA as a potently neuroactive compound from the tunicate, *Didemnum* sp. (**Figure 1**). The tunicate was identified to genus by sequencing the mitochondrial cytochrome *c* oxidase I (COXI) gene (GenBank MW728885). To determine DIMTA’s mechanism of action, we monitored its effects on mouse DRG neurons using constellation pharmacology. The neurons were stimulated with 25 mM KCl, which elicited calcium signals. Subsequently, the cells were incubated with DIMTA for six minutes, and the effects of the compound were assessed on basal intracellular calcium levels and on the signal elicited by the subsequent application of KCl. At a concentration of 25 µM, DIMTA elicited two notable effects on LTCT: the baseline calcium levels were elevated during the incubation period of the compound, referred to henceforth as direct effects of the compound; and the compound suppressed the magnitude of calcium signals elicited by the subsequent application of 25 mM KCl, henceforth referred to as indirect effects. In other types of DRG neurons, DIMTA (25 µM) only exhibited indirect effects. DIMTA had no visible effects on glia at 2.5 or 25 µM. These data suggested that at a high concentration of 25 µM, the compound hit two different molecular targets: one that resulted in the elevation of baseline calcium levels and the other that suppressed the depolarization-induced calcium influx. To identify the high-affinity molecular target, we focused on the specific subpopulation of neurons that were affected by DIMTA at lower concentrations. At a lower DIMTA concentration (2.5 µM) (tested in n=3 experiments, monitoring an average of 422 neurons and 602 glia from mouse DRG in each experiment), we found that the direct effects were observed solely in LTCTs: 64% (± 7%) of LTCTs exhibited an increased baseline in response to DIMTA application, indicating that more cytoplasmic Ca^2+^ was present in the cells in the presence of DIMTA, while no other cell type exhibited direct effects (**Figure S1**). In contrast, DIMTA’s indirect effects at low concentrations were greatly diminished: only 16% (± 9%) of all neurons were partially inhibited (**Figure S1**).

**Figure 1.**
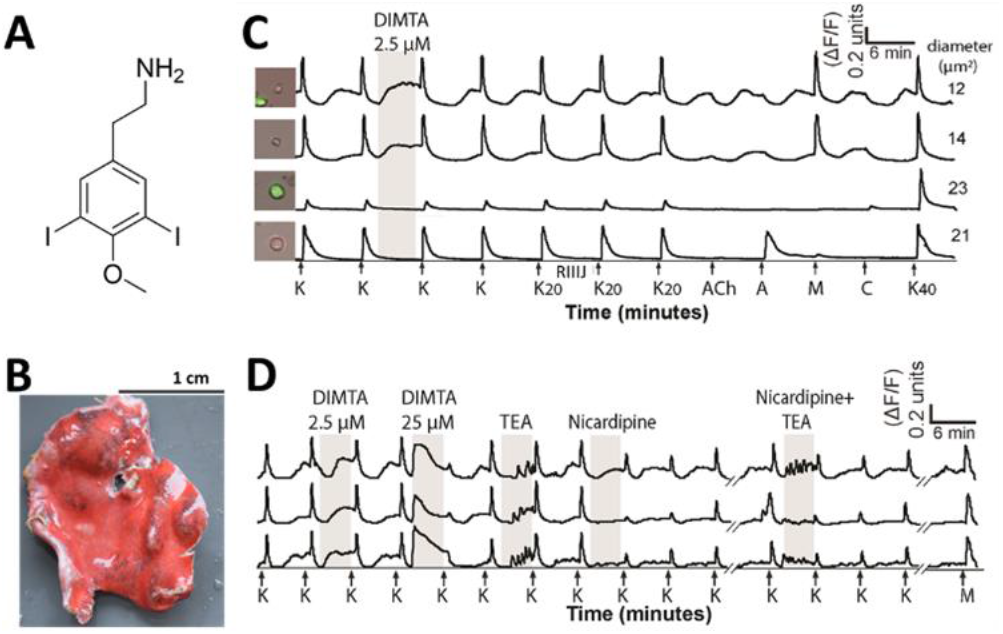
Constellation pharmacology shows the effects of DIMTA on different subsets of DRG neurons. **(A)** DIMTA was isolated from (**B**) the tunicate *Didemnum* sp. (SI-028S). **(C)** and (**D**) represent constellation pharmacology using DIMTA. Each trace follows the Fura2-AM fluorescence ratio at 340/380 nm (a measure of relative intracellular calcium levels) (*y*-axis) of a single cell out of ∼500 neurons in a single experiment. Chemicals were added over the experiment time (*x*-axis), and the cellular responses were recorded. Arrows indicate 15-s application of potassium chloride (K, 25 mM or K_20_, 20 mM). Sequential application of a set of pharmacological ligands was used to identify neuronal cell populations: K_20_, κM-RIIIJ (1 µM), acetylcholine (ACh, 1 mM), allyl isothiocyanate (A, 100 µM), menthol (M, 400 µM), and capsaicin (C, 300 nM). The column on the left shows the bright-field image of cells from which the calcium traces were obtained. The highlighted region indicates the period of DIMTA application. Potassium chloride 40 mM (K_40_) was applied at the end of the experiment to determine the viability of the neurons. (**C**) The top two traces are LTCTs that displayed a direct effect to treatment with DIMTA (2.5 µM). The bottom two traces are representative of neurons that were not affected by the compound. **(D)** DIMTA exhibits features of VGKC and VGCC block in constellation pharmacology. Selected traces from LTCTs illustrating the effects of DIMTA at 2.5 µM and 25 µM. TEA (10 mM) is a wide spectrum K_v_ channel antagonist, while nicardipine (400 nM), is a Ca_v_1 channel antagonist.

In this assay, LTCTs are characterized pharmacologically in part by their response to menthol, but not to allyl isothiocyanate (AITC), reflecting the importance of the cold-sensing menthol receptor transient receptor potential M8 (TRPM8)^15^ and the absence of the nociceptive transient preceptor potential A1 (TRPA1) in LTCTs. Furthermore, these cells have a characteristic calcium profile identified by the distinct transient decrease or dip in cytosolic calcium concentration [Ca^2+^]_i_, a fluctuating baseline that reflects their responsiveness to very small drops in temperature of <1 °C.^6^ DIMTA increased the baseline [Ca^2+^]_i_ of LTCTs, suggesting that targeting DIMTA’s high affinity molecular target modulates the temperature of activation in thermosensors.

### DIMTA blocks voltage-gated ion channels in LTCTs

DIMTA (25 µM) exhibited paradoxical effects on LTCTs: it elevated the baseline (increased influx of calcium), yet inhibited the response of the cells to 25 mM KCl (decreased the influx of calcium) (**Figure 1**). Previously, it was observed that, the promiscuous K^+^-channel blocker tetraethylammonium chloride (TEA) at 10 mM concentrations elevated the baseline of LTCTs, causing repetitive [Ca^2+^]_i_ spikes.^4, 16^ In addition, because LTCTs predominantly express Ca_v_1 channels, application of the Ca_v_1 antagonist nicardipine attenuates the responses of these neurons to K^+^-induced depolarization.^6^ Thus, we hypothesized that co-application of TEA and nicardipine would replicate the effects of DIMTA (25 µM) on DRG neurons.

TEA (10 mM) alone caused elevated spiky [Ca^2+^]_i_ responses to the baseline, while nicardipine (400 nM) alone partially blocked K^+^-elicited calcium responses (Figure 1). When TEA and nicardipine were co-applied, the LTCTs exhibited a combination of both responses that were similar, but not identical, to the phenotypic effects observed from the application of DIMTA (25 µM). As found with DIMTA, the co-application of TEA and nicardipine led to both an elevated baseline and a blocked response to KCl in LTCTs. However, the increased baseline in TEA exhibited a spiky shape, while the DIMTA-increased baseline was very smooth. This shape difference had two potential causes: DIMTA and TEA target different K_v_ channel subtypes; alternatively, the resolution of capturing the calcium signals (frame rate of every 2 seconds) may be sufficient to capture calcium spikes elicited by TEA but insufficient to capture spikes from the application of DIMTA. Overall, this comparison suggested that the observed phenotypic effects of DIMTA may originate from targeting two different classes of voltage gated ion channels (VGICs): voltage gated potassium channels (VGKCs) and voltage gated calcium channels (VGCCs).

Bay K 8644 is a dihydropyridine that directly interacts with VGCCs to induce calcium entry, instead of the VGCC block induced by most dihydropyridine drugs. Thus, co-application of Bay K 8644 with KCl should induce additional depolarization in DRGs due to increased calcium entry. In support of the hypothesis that DIMTA affects VGCCs at 25 µM, DIMTA blocked the additional depolarization induced by the co-application of Bay K 8644 (200 nM) and KCl (20 mM) (**Figure S2A**).

To determine whether DIMTA also affects voltagegated sodium channels (VGSCs), we applied the VGSC agonist *Anemonia viridis* toxin 2 (ATX-II; 100 nM) which delays sodium channel inactivation. We then tested the effect of applying DIMTA (2.5 µM, 25 µM) and the sodium channel blocker tetrodotoxin (TTX; 1 µM) before and after ATX-II application. DIMTA’s direct effect on LTCT depolarization did not change significantly in the presence of ATX-II (**Figure S2B**). These results suggest that the direct effect of DIMTA does not rely on VGSCs, or that its impacts on VGSCs are not observable in DRG neurons.

### DIMTA blocks voltage-gated ion channels in LTCTs

DIMTA (2.5 µM) was applied to DRG neurons, leading to identification of LTCTs that exhibited an increased baseline in response to DIMTA. Single cells with this phenotype were selected for tandem whole-cell electrophysiological experiments to determine the effects of compound on the firing properties of the cell (**Figure 2A**). We monitored the membrane potential of the cell at resting condition (no current injected, **Figure 2B**). As an internal control, the bath solution without DIMTA was applied, leading to no change of the resting potential. By contrast, application of DIMTA (2.5 µM) elicited firing of action potentials. The effects of DIMTA were slowly reversible, as observed by the sustenance of action potential firing even after the washout of the compound. Additionally, we monitored the effects of the compound on action potential profiles by using current injection pulse protocols. We maintained the membrane potential of the cell at ∼-70mV by injecting current, and compared the effects of DIMTA (2.5 µM) after injecting pulses of negative and positive currents for 500 ms. DIMTA reduced the magnitude of afterhyperpolarization (AHP) by 8 ± 3.5 mV (n=11 LTCT cells, p<0.001) (**Figure 2C**), resulting in depolarization induced inactivation at current injections greater than 100 pA (**Figure 2D**).

**Figure 2.**
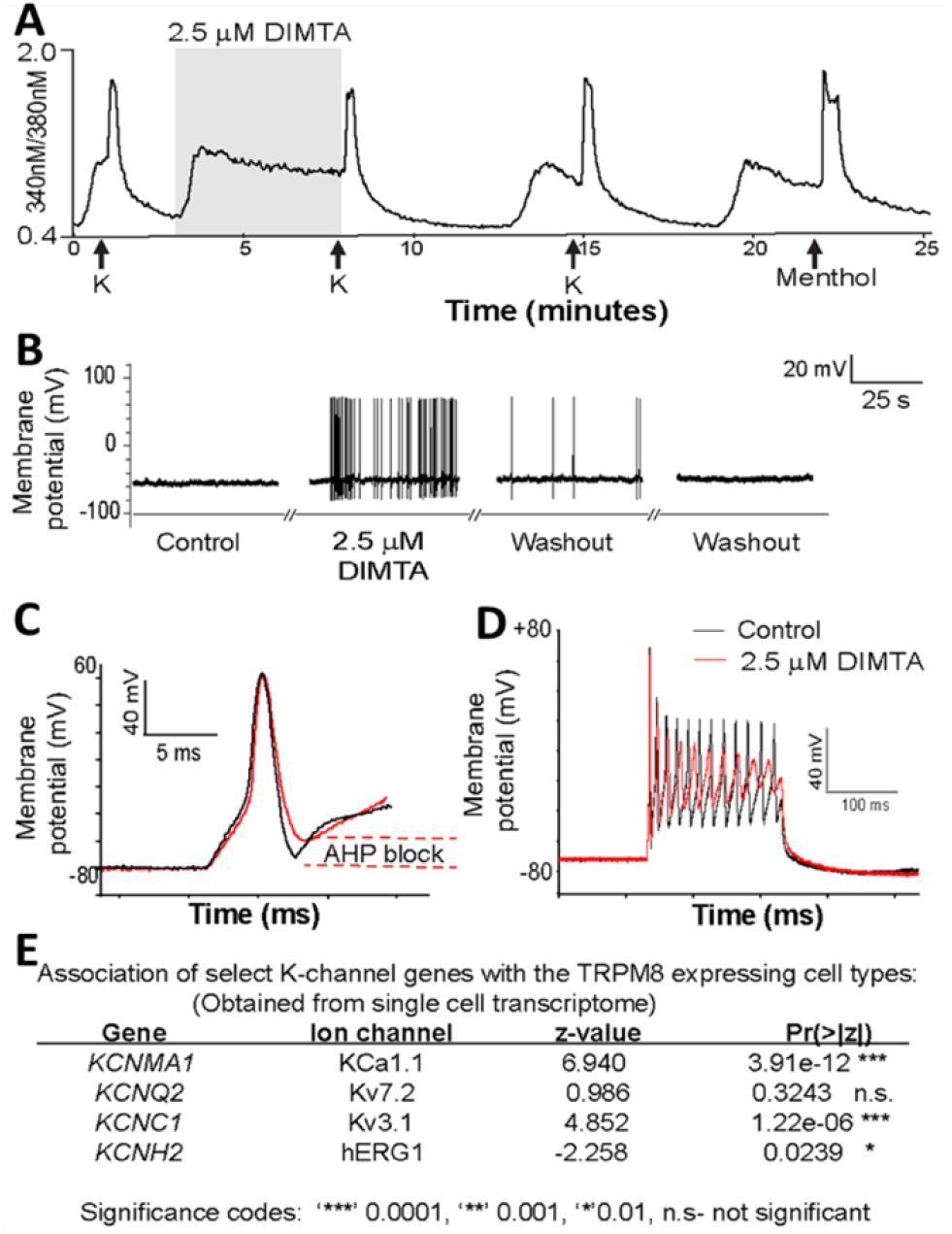
Identifying the molecular target of DIMTA in LTCTs by constellation pharmacology in tandem with single-cell electrophysiology and transcriptomics analysis. **(A)** Calcium imaging trace of an LTCT with an increased baseline in response to DIMTA (2.5 µM). The axes and abbreviations are the same as described in Figure 1. **(B)** Tandem whole-cell electrophysiology experiment performed on the same LTCT neuron indicating that application of DIMTA (2.5 µM) caused firing of action potentials. **(C)** Monitoring the effects of DIMTA on action potential profiles with current injection pulse protocols. DIMTA reduced the magnitude of the afterhyperpolarization (AHP) by 8 ± 3.5 mV (n=11 LTCT cells, p<0.001). **(D)** Depolarization induced inactivation at current injections greater than 100 pA with the reduction of AHP magnitude. **(E)** Association of K-channel genes with TRPM8 expressing cell types.

AHP is modulated by VGKCs including K_V_7.2 and by calcium-activated K_Ca_3. Using previously published single-cell RNA sequencing (scRNA-seq) data,^17^ we identified several potassium channels that are strongly expressed in TRPM8-expressing DRG neurons (**Table S1**). We also selected neurons identified as LTCTs by constellation pharmacology and sequenced those cells. Channels that were expressed in both data sets included K_Ca_1.1 and K_V_3.1 (**Figure 2E**). Transcripts for Kv7.2 were also observed in these cells but were not significantly correlated with this cell type, in comparison to others. We therefore proposed that the AHP induced by DIMTA might result at least in part by blocking K_V_7.2, K_V_3.1, and/or K_Ca_1.1, as well as other potassium channels expressed in LTCT neurons.

### DIMTA is a modest inhibitor of K_Ca_1.1, K_v_3.1, and K_v_11.1 but does not block K_v_7.2 or TRPM8

Using expressed human K_V_7.2, K_V_3.1 and K_Ca_1.1 in *Xenopus laevis* oocytes, we performed two-electrode voltage-clamp recordings (**Figure 3**). Applying voltage step protocols, we elicited currents in the absence and presence of DIMTA. DIMTA (5 µM) inhibited K_Ca_1.1 (18 ± 5%, n=5) and K_V_3.1 (25 ± 5%, n=5), but did not affect K_V_7.2.

**Figure 3.**
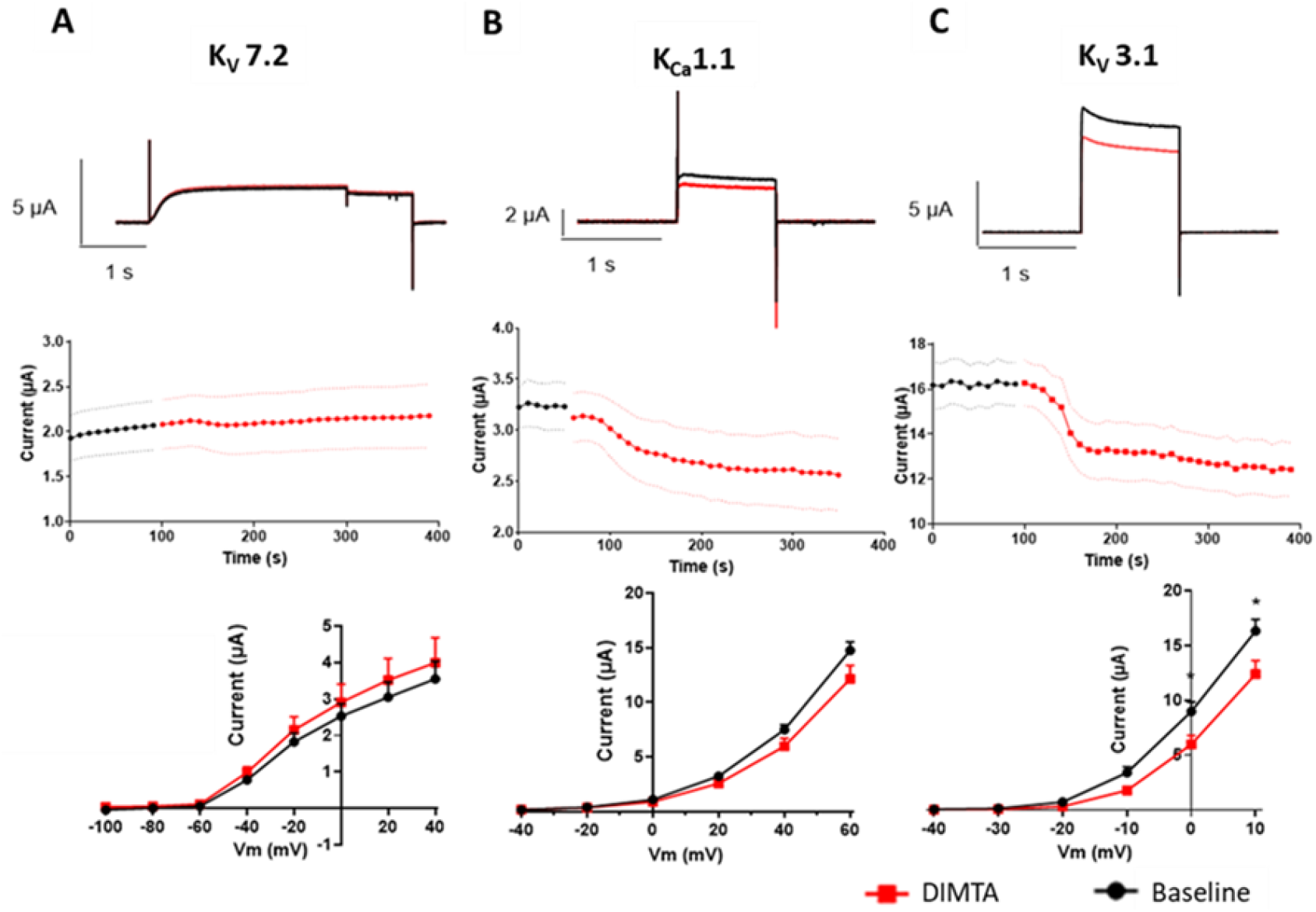
Effects of DIMTA on *Xenopus laevis* oocytes expressing **(A)** K_V_7.2, **(B)** K_Ca_ 1.1, and **(C)** K_V_3.1. Top: Representative current traces before (black) and after (red) application of DIMTA (5 µM). Middle: Mean current-time plots. Bottom: Current-voltage relationships in the absence and presence of DIMTA (5 µM). Data are presented as mean ± SEM, n=5.

We performed patch-clamp experiments using HEK-293 cells constitutively expressing human K_Ca_1.1 (**Figure 4**). As expected, depolarizing the cells in the presence of 100 nM free intracellular calcium gave rise to outward potassium currents (**Figure 4A**). Extracellular application of increasing concentrations of DIMTA caused a concentration-dependent inhibition of the hK_Ca_1.1 current at all activating membrane potentials with an IC_50_ value of 16 ± 2 µM, whereas control runs did not decrease current (**Figure 4B-4C**). Subsequent perfusion with compound-free extra-cellular solution reversed the inhibitory effect. By analyzing the G/Gmax curve, we also found that DIMTA did not change the voltage dependence of activation (**Figure 4D**).

**Figure 4.**
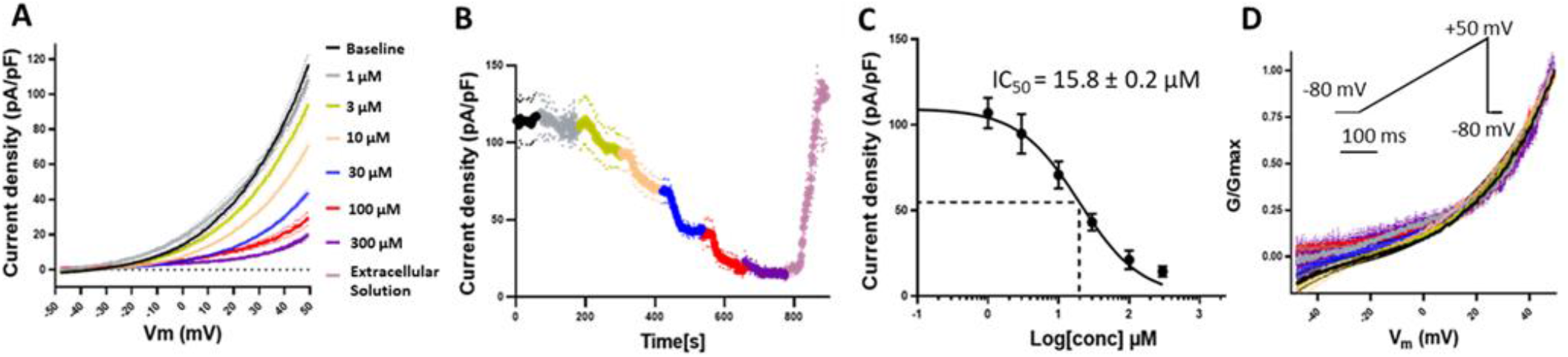
Effect of DIMTA on hK_Ca_1.1channels expressed in HEK-293 cells. **(A)** Mean current-voltage relationships and **(B)** current-time plots before and after application of increasing concentrations of DIMTA, n=6-13. **(C)** Concentration-response curve summarizing the effect of increasing concentration of DIMTA on hK_Ca_1.1. Currents were measured at +50 mV (n=13). **(D)** Conductance curves (G/Gmax curve) (n=9-13). Data are presented as mean ± SEM. Currents were elicited every 2 s using the voltage-clamp protocol shown in the inset.

We conducted inside-out membrane patch recordings to determine whether DIMTA also blocks from the intracellular side of the membrane (**Figure S3**). We observed that DIMTA (30 μM) inhibited the K_Ca_1.1 current to a comparable extent both inside-out and extracellular membranes (50% inhibition vs. 62%, respectively). However, the time course of inhibition was slower when DIMTA was applied to the inside than when it was applied to the outside. We did not observe any change in time matched control experiments (**Figure S4**).

DIMTA reversibly inhibited K_V_3.1 channels in transiently transfected HEK-293 cells in a concentration dependent manner (IC_50_ 11.8 ± 0.2 µM; **Figure 5**). Although we did not observe changes of the voltage dependence of activation, we found that DIMTA affected inactivation, shifting the voltage dependence of inactivation to more negative potentials (**Figure 6)**.

**Figure 5.**
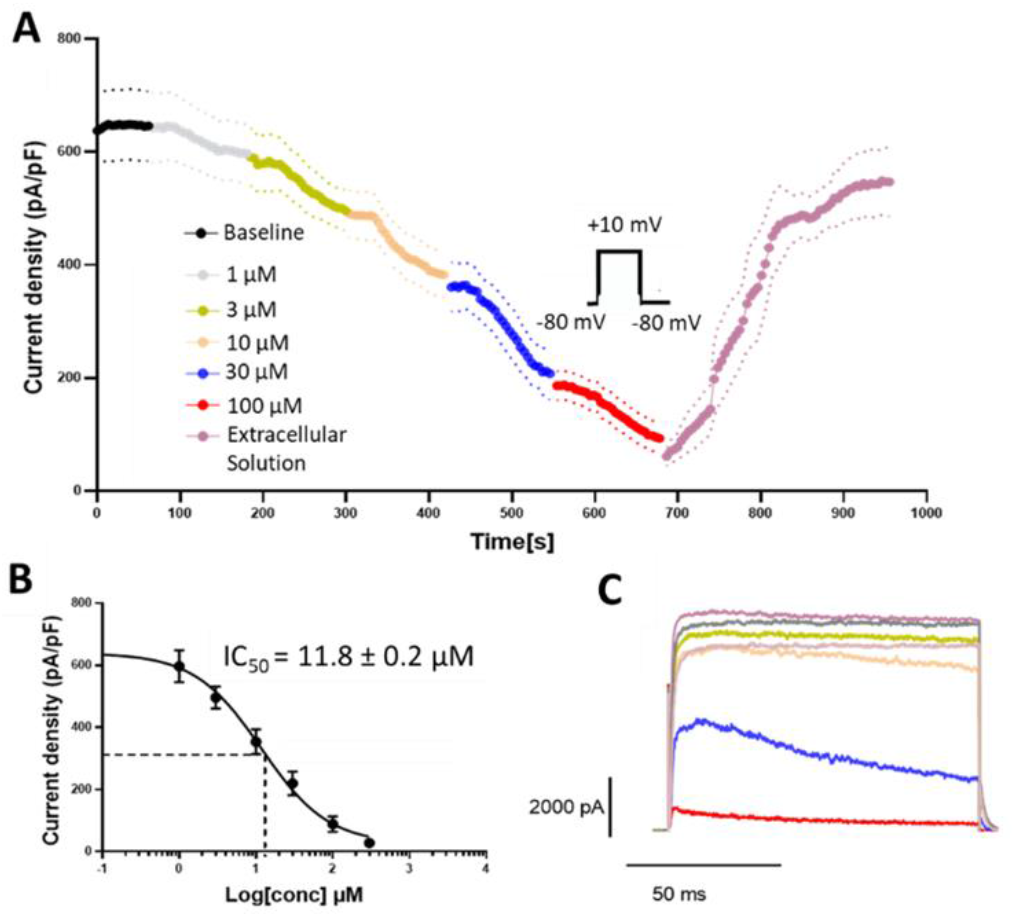
Effects of different concentrations of DIMTA on K_V_3.1 expressed in HEK-293 cells. **(A)** Current time plot of the effects of 1 to 100 µM DIMTA (n=6-18). **(B)** Concentration response relationship for DIMTA (n=18). To get the full inhibition of the K_V_3.1 current additional experiments with only 300 µM DIMTA were performed (n=7). **(C)** Representative current recordings before and after application of increasing concentrations of DIMTA. Currents were elicited with the voltage protocol shown in **(A)**. Data are presented as mean ± SEM.

**Figure 6.**
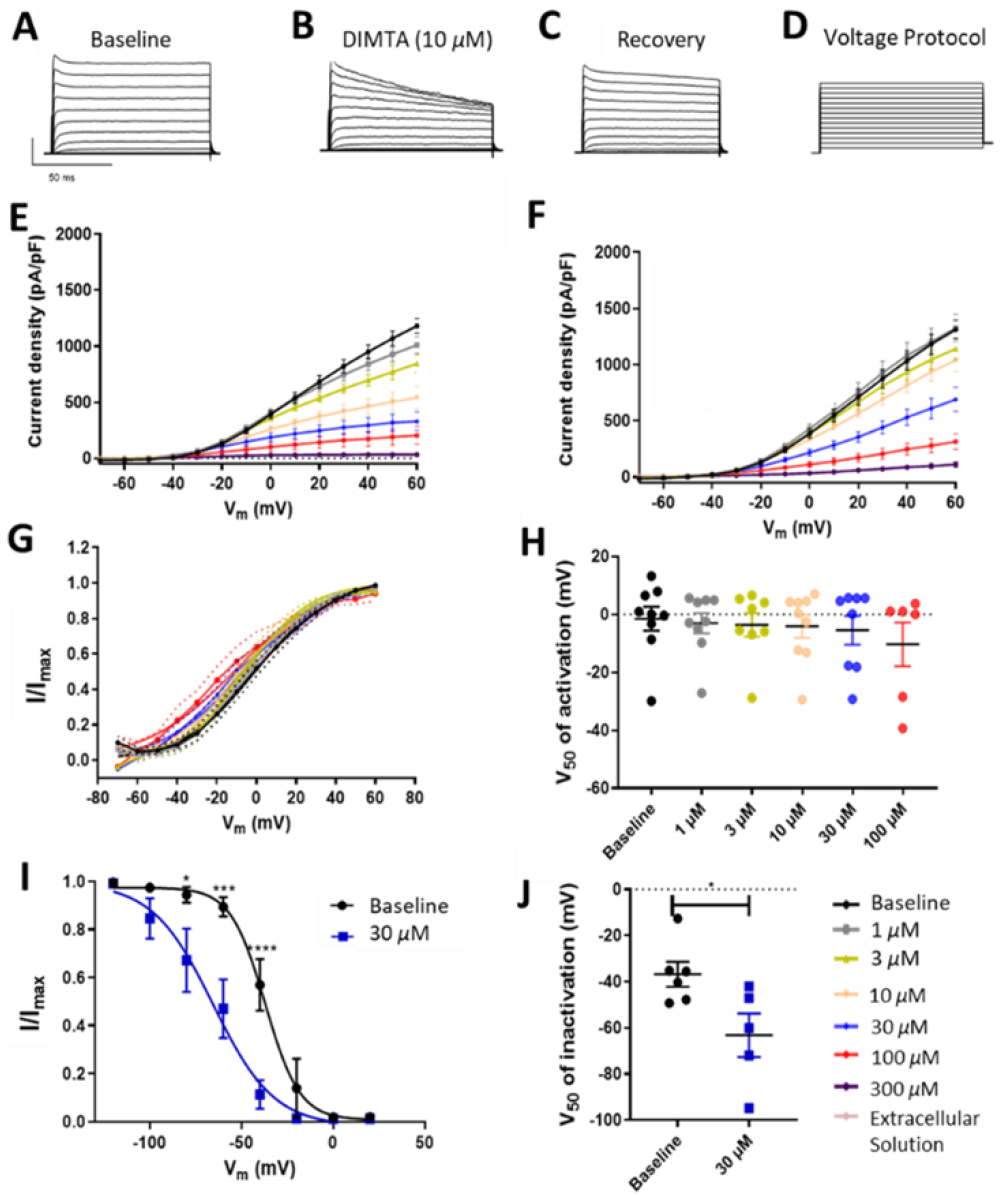
Effects of DIMTA on K_V_3.1 expressed in HEK-293 cells. **(A)** Examples of current traces recorded at baseline; **(B)** In the presence of DIMTA (10 µM); **(C)** Upon washout of the compound. Currents were elicited using the voltage protocol shown in **(D)**. Current-voltage relationship before and after increasing concentrations DIMTA when constructed by plotting the steady state current **(E)** or peak current **(F)** as a function of membrane potential, n=6-19. (G) Voltage dependence of channel activation, n=7-14. **(H)** Membrane potential of half-maximal activation (V_50_). **(I)** Steady state inactivation curves. **(J)** V_50_ of inactivation, n=6-9. Data are presented as mean ± SEM.

K_v_11.1 (also known as hERG) is a potassium-selective ion channel essential for repolarization of the cardiac action potential. While other potassium channels were tested because they affect AHP, K_V_11.1 was tested because it is considered crucial in drug discovery programs, since it prolongs the QT-interval in the electrocardiogram and increases the risk of ventricular tachycardia.^18^ Using Chinese Hamster Ovary (CHO) cells constitutively expressing K_V_11.1 channels (**Figure S5**), we found that DIMTA inhibits K_V_11.1 (IC_50_ 18.7 ± 0.1 µM).

The data show that DIMTA is a VGKC inhibitor. Modest effects on AHP are accounted for by inhibition of K_V_3.1 and K_Ca_1.1. We speculate that the baseline shift seen in LTCTs is caused by a more potent inhibition of other K_V_ channels expressed by LTCTs (**Table S1**), although the potentially enormous number of heteromeric K-channels in LTCTs makes this impossible to fully evaluate using current technology. To evaluate whether DIMTA targeted cold sensor TRPM8, we analyzed TRPM8-overexpressing (hTRPM8) HEK-293 cells in several conditions. Inhibition was only detected at very high concentration of DIMTA (IC_50_ = 112 µM), indicating that TRPM8 is not a relevant target of DIMTA (**Figure S6**).

### DIMTA dampens cold sensitivity

4-Aminopyridine (4-AP) is an FDA-approved drug for multiple sclerosis that broadly blocks K^+^ channels, albeit at a >10-fold higher concentration than DIMTA, and it inhibits calcium and other channels at even higher concentrations.^19^ 4-AP makes mice more sensitive to temperature changes.^20^ Since we anticipated that DIMTA operates similarly, we performed behavioral studies in mice to test its effect on the latency and temperature threshold for eliciting a cold pain withdrawal response. In blinded experiments, mice were injected with vehicle (1% DMSO in saline) or 1 mg/kg or 10 mg/kg DIMTA. After 30 min, animals were placed on cold plate held at room temperature for 3 min, and subsequently the temperature of the plate was ramped down at the rate of 10 °C/min. The time to respond to decreasing temperature was measured, and the corresponding temperature at first response was noted (**Figure 7**). Vehicle-injected control animals withdrew their paws after 96 s and at 8 °C, while animals injected with 10 mg/kg DIMTA moved their paws after 122 s (\**p* < 0.05) and at 4.6 °C (*p* = 0.06). Behavioral response with DIMTA at 1 mg/kg was not significantly different from the vehicle (*p* > 0.05). Thus, in contrast to our hypothesis, DIMTA at 10 mg/kg significantly increased the threshold of cold temperature sensitivity. This effect might be due to the diminished AHP observed in LTCTs treated with DIMTA, which results in faster inactivation of Na^+^ channels, thereby causing desensitization of these cell types.

**Figure 7.**
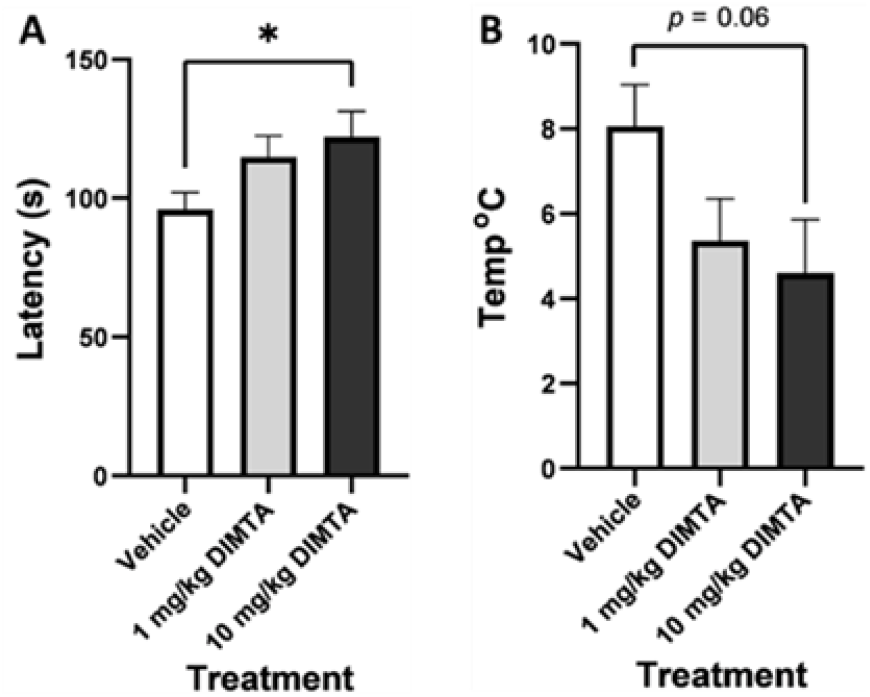
Male adult (6-8 weeks old) CD1 mice were injected *i*.*p*. with DIMTA (1 mg/kg, 10 mg/kg), or vehicle (1% DMSO in saline) 30 min prior to behavioral testing. Mice were assessed in a cold-plate chamber in which the temperature was decreased from room temperature at a rate of 10 °C per min. Pain-related behaviors including lifting and licking of the forepaw were observed. The **(A)** time and **(B)** temperature of the first sign of pain were recorded. Values shown by the histograms are expressed as the mean ± standard error of the mean (n = 13). The compound effect versus vehicle (\**p*<0.05). Statistics were evaluated by one-way Anova and Dunnett’s multiple comparisons test; the difference in values between panels A and B results from the higher precision level used in time measurement versus temperature measurement.

## CONCLUSION

One of the crucial goals in discovering effective drug leads is to identify target-selective compounds. However, when disease relevant molecular targets are widely expressed in many cell types, it is difficult to predict *a priori* how a target-selective drug will impact complex mammalian physiology. In addition, it is challenging to assess the target selectivity of new compounds on complex targets such as the potassium channels. To circumvent these problems, a shift in drug discovery paradigm is required that focuses on the identification of ligands that selectively target specific neuronal cell types, rather than specific molecular targets. In this work, we describe a strategy to achieve this goal.

The sensory neurons of the DRG represent many cell types that are specialized to relay specific sensory information to the central nervous system. Previous work has defined 16 specific cell types in the DRGs, used to screen ligands for the management of neuropathic pain.^4^ Here we identified and characterized one of the bioactive ligands from our library of marine natural products, DIMTA (**Figure 1**). The compound was active in constellation pharmacology assays performed on dissociated neurons from the mouse DRG, selectively targeting just one of the cell types (LTCTs) at lower doses (**Figure 1**). To identify the molecular target underlying DIMTA’s cell-type selectivity, we performed tandem single-cell electrophysiology and single-cell transcriptomic analyses on LTCTs. As we demonstrated in **Figure 2**, DIMTA blocked AHP in LTCTs; subsequently a list of putative molecular targets that contribute to AHP in LTCTs were identified from single-cell transcriptomics. By comparing the expression of these molecular targets with the other unaffected cell types in the primary culture, we narrowed the target to three K-channels that might, in part, underlie AHP phase of action potentials in LTCTs: K_V_7.2, K_V_3.1, and/or K_Ca_1.1 (**Figure 2D**). These molecular targets were further tested and validated using a heterologous expression system, and DIMTA was found to block K-channels KCa1.1, Kv 3.1 and Kv11.1, but not target Kv7.2. Furthermore, our experiments indicate that the drug acts as pore blocker (there is no change in the voltage dependence of activation, and it works from both sides of the membrane).

To assess the physiological effects of DIMTA *in vivo*, particularly its effects on cold sensation, we tested DIMTA using cold plate assays.^21^ Our data suggests that the administration of DIMTA attenuates cold response behaviors. However, full elucidation of this effect will require optimized ligands with higher target selectivity. Since DIMTA is chemically simple, it provides an attractive target for such optimization. Cold allodynia is one of the most commonly experienced chronic pain symptoms in a range of neuropathic pain syndromes caused by nerve injury, tissue damage, and chemotherapy.^22, 23^ The compound described in this work represents a scaffold that can be potentially optimized by structure-activity relationship studies for useful therapeutics to alleviate pain.

## MATERIALS AND METHODS

### Animals

The use of all animals followed protocols that were approved by the Institutional Animal Care and Use Committee of the University of Utah. To identify somatosensory neuronal subclasses in the calcium imaging experiments, calcitonin gene-related peptide (CGRP)-green fluorescent protein (GFP) transgenic reporter mice were used. Strain STOCK Tg(Calca-EGFP)FG104Gsat/Mmucd) was created by the Gensat project as previously described.^24^ In this mouse strain, GFP expression is driven by the gene regulatory elements of CGRP, which primarily labels peptidergic nociceptors in the somatosensory neuronal cell population. Male CD1 mice (20-30 g, 6-8 weeks old) obtained from Charles River Laboratories were acclimated for at least 5 days before the start of treatment.

### Cell culture

DRG neurons harvested from CGRP-GFP mice were cultured in minimum essential medium (MEM, pH = 7.4) supplemented with 10% fetal bovine serum (FBS), penicillin (100 U/mL), streptomycin (100 µg/mL), HEPES (10 mM), and glucose (0.4% w/v). Human embryonic kidney 293 (HEK-293) cells (ATCC) stably expressing human TRPM8 were cultured in Dulbecco’s modified eagle’s medium: Nutrient Mixture F12 Ham (DMEM:F12) media supplemented with 5% FBS and Geneticin (300 µg/mL). HEK-293 and Chinese hamster ovary (CHO) cells were cultured in DMEM supplemented with 10% FBS and 1% penicillin-streptomycin. All cells were cultured in an incubator maintained at 37 °C in a humidified atmosphere with 5% CO_2._

### Expression constructs

Plasmids suitable for heterologous expression in *Xenopus laevis* oocytes and mammalian cells harboring the cDNAs of human K_V_7.2 (*KCNQ2*, GenBank NM_004518) and K_Ca_1.1. (*KCNMA*, GenBank NM_002247) have been described previously.^25, 26^ The plasmid carrying human K_V_3.1 (*KCNC1*, GenBank NM_001112741.1)^27^ was a kind gift from Dr. Thomas Jespersen (University of Copenhagen, Denmark).

### General chemical procedures

NMR data were collected using a Varian INOVA 500 spectrometer operating at 500 MHz for ^1^H and 125 MHz for ^13^C, and equipped with 5 mm Varian HCN Oneprobe for proton detected experiments and a 3 mm Varian inverse probe for carbon detected experiments. NMR shift values were referenced to the residual solvent signals. UPLC-HRESI(+)-TOFMS analysis was performed on a Waters Acquity H class UPLC system coupled to a Waters Xevo G2-XS qTOF equipped with a Z spray ESI source. HPLC separations were performed using a Thermo Scientific™ Dionex™ Ultimate-3000 HPLC system equipped with a photodiode array detector. The purity of DIMTA was rigorously determined to be >99% using UPLC-MS, HPLC-DAD, and NMR experiments. The amount of DIMTA used in pharmacology experiments was measured with high accuracy using a digital analytical balance (Mettler Toledo).

### Collection, extraction of genomic DNA, and sequencing of phylogenetic markers

The tunicate *Didemnum* sp. was collected by hand using SCUBA in April 2018 (collection number SI-028S) from Solomon Islands (S 09° 6’51.11 E 160°11’2.10). Portions of the freshly collected sample were set aside for chemical analysis and preserved in RNAlater and were kept frozen at −20 °C until use. Genomic DNA was extracted from the RNAlater-preserved tissue of SI-028S using Genomic-tip (Qiagen). The mitochondrial COX1 gene was amplified from the genomic DNA using primers LCO1490 (5’-GGT CAA CAA ATC ATA AAG ATA TTG G) and HCO2198 (5’-TAA ACT TCA GGG TGA CCA AAA AAT CA) ^28^. The polymerase chain reaction was performed using a 50 µL master mix consisting of 1x standard *Taq* buffer (New England Biolabs), LCO1490 primer (0.2 mM), HCO2198 primer (0.2 mM), dNTP mix (200 µM), *Taq* DNA Polymerase (1.25 U, New England Biolabs) and template DNA (10 ng/µL). PCR conditions were as follows: a hot start (94 °C, 2 min) followed by 39 cycles of [94 °C/30 s, 45 °C/30 s, 72 °C/2 min], then a final extension at 72 °C for 10 min. PCR product was purified using the QIAquick Gel Extraction Kit (Qiagen) and Sanger sequenced (Genewiz).

### Isolation and purification of DIMTA

Frozen sample of SI-028S (wet weight, 50 g) thawed, diced, and exhaustively extracted with CH_3_OH. The resulting extract was filtered, dried *in vacuo* and subjected to reversed-phase HPLC using a Thermo Scientific™ Dionex™ WPS-3000 HPLC system equipped with a Photodiode array detector. The sample was purified using a Phenomenex Luna C_18_ column (250 x 10 mm), eluting with an isocratic condition consisting of 30% CH_3_CN in H_2_O (0.1% TFA) over 20 min at 3.5 mL/min flow rate to yield a white amorphous powder identified as DIMTA (31 mg) by comparison of its ^1^H NMR, ^13^C NMR, and HRMS data with those previously reported (**Figure S7-S8**).^10, 11^ The HPLC-purified compound (**Figure S9**) was dissolved in DMSO to obtain concentrated stock solutions (12.5 mM) that were maintained frozen (−20 °C). The stock solutions were thawed and diluted to their final concentrations on the day of the experiments.

### DRG neuron preparation and calcium imaging

DRG neuron protocols have been described in detail previously.^4, 29-31^ Briefly, lumbar DRG neurons were harvested from CGRP-GFP mice in a CD-1 genetic background. The DRG neurons were dissociated by trypsinization and mechanical trituration and were subsequently plated onto 24-well poly-D-lysine-coated plates. Lastly, DRG neurons were cultured and incubated overnight at 37 °C, 5% CO_2_ in MEM.

The cultured DRG neurons were incubated with Fura2-AM (2.5 µM) (Molecular Probes) in MEM at 37 °C for 1 h, and at r.t. for 0.5 h prior to imaging. The MEM solution was then replaced at least twice with the observation solution (NaCl (145 mM), KCl (5 mM), CaCl_2_ (2 mM), MgCl_2_ (1 mM), sodium citrate (1 mM), HEPES (10 mM), glucose (10 mM)). The relative level of intracellular Ca^2+^ concentration upon depolarization with 25 mM KCl was monitored by measuring the changes in emitted fluorescence at 510 nm from 340 nm/380 nm excitation signals. To assess the effects of DIMTA on DRG neurons, KCl solution was applied for 15 s at regular intervals of 7 min and after the second depolarizing pulse, the compound was incubated with the cells for 5 min. Following incubation, a third depolarizing KCl pulse was applied to determine the effect of DIMTA on the responses of the neurons to depolarization. Succeeding depolarizing pulses were done to determine the reversibility of the observed responses of the neurons to depolarization after compound incubation. Each experiment was followed by sequential application of a set of pharmacological challenges to identify neuronal cell populations. Pharmacological challenges present in each experiment include K^+^ (20 mM and 40 mM), κM-RIIIJ (1 µM), allyl isothiocyanate (AITC; 100 µM), menthol (400 µM), and capsaicin (300 nM). After each calcium imaging experiment, the cells were incubated with Hoechst stain (1000 µg/mL) for 5 min at rt and then washed 3 times. Subsequently, the cells were incubated with Alexa-Fluor 647 Isolectin (IB4, 2.5 µg/mL) for 5 min at r.t. and then washed 3 times. Cell images were acquired using a rhodamine filter set. Nis elements and CellProfiler^32^ were used to acquire and create ROIs and extract cellular information, respectively. Video information and trace data were extracted using an in-house script built in Python and R language.

### Electrophysiological recordings from cultured DRG neurons

Whole cell current clamp and voltage clamp experiments were performed in tandem with calcium imaging. LTCTs were identified by calcium imaging experiments and selected for whole cell recordings. For tandem recordings, the intracellular pipette solution contained: potassium aspartate (140 mM), NaCl (13.5 mM), MgCl_2_ (1.8 mM), EGTA (0.09 mM), HEPES (9 mM), creatine phosphate (14 mM), Mg-ATP (4 mM) and Tris-buffered GTP (0.3 mM). The pH of the solution was adjusted to 7.2 with KOH, and the osmolarity was adjusted to 290-300 mOsM with glucose. The extracellular bath solution was the same as the DRG observation solution used for calcium imaging experiments and contained: NaCl (145 mM), KCl (5 mM), CaCl_2_ (2 mM), MgCl_2_ (1 mM), sodium citrate (1 mM), HEPES (10 mM), and glucose (10 mM). The pH of extracellular solution was adjusted to 7.4 with NaOH, and the osmolarity was adjusted to 310-320 mOsM with glucose. The pipette resistance ranged from 3-5 MΩ for voltage clamp experiments and 5-10 MΩ for current clamp experiments. Cells with stable resting membrane potential below −40 mV were used for recordings, which were made with a MultiClamp 700A amplifier and acquired with a DigiData 1440 digitizer. The amplifier and digitizer were under controlled by MultiClamp Commander and Clampex10.6 (Molecular Devices), respectively. All experiments were done at room temperature (22 °C).

To test the effects of the compound (DIMTA) on membrane potential, the cells were patched and membrane potential was monitored at rest (no current injections). The bath solution (control) was applied followed by the application of the DIMTA. The cells were washed and a wait time of 10 min was given to allow recovery from the effects of the compound. A pulse protocol was then designed to monitor the effects of DIMTA on action potential. Cells were held at resting potential (0 pA current injection) followed by a 500 ms pulse of varied current injections (- 125 pA to +150 pA current in 25 pA steps). The same protocol in the presence of DIMTA, after washout of DIMTA, and in a control containing the bath solution in place of DIMTA was performed to compare the effects of the compound.

To monitor the effects of DIMTA on whole cell currents, the cells were held at −70 mV and steps of depolarizing stimulus were given (−120 mV and −60 mV to + 40 mV in steps of 10 mV). The whole cell inward and outward currents were recorded at ∼70% series resistance compensation. The same protocol was run in the presence of the DIMTA, after washout of DIMTA, and in a control containing the bath solution in place of DIMTA.

### Single-cell transcriptomics analysis

As previously described,^4^ individual cells were picked using fire-polished glass pipettes with optimized diameter after completion of the constellation pharmacology experiments. Cells were lysed and mRNA was reverse transcribed to generate cDNA, which then underwent whole transcriptome amplification, all using the QIAseq FX Single Cell RNA library kit according to the manufacturer’s standard protocol (Qiagen). The amplified cDNA was used to construct a sequencing library for the Illumina NGS platform, also using the QIAseq FX Single Cell RNA library kit. The amplified cDNA was fragmented to 300 bp in size, treated for end repair and A-addition, followed by adapter ligation and then cleanup with Agencourt AMPure XP magnetic beads (Beckman Coulter Life Sciences). The cDNA library was submitted to the High Throughput Genomics Shared Resource, Huntsman Cancer Institute, for library quality control and sequencing. Sequencing data was analyzed using in house R scripts described before.^4^

### Fluorometric calcium flux assay with HEK-293 TRPM8 overexpressing cells

Prior to calcium imaging experiments, the cells were seeded on a 96-well plate pre-coated with 1% gelatin, grown to confluence and incubated with the calcium indicator Fluo 4-AM (Fluo-4 Direct assay kit, Invitrogen) for 60 min at 37 °C. The loading solution was replaced with a wash solution comprising of LHC9 media (Life Technologies), 1 mM probenecid, and 0.75 mM trypan red (ATT Bioquest) 30 min before activity analysis. For inhibition assays the samples were supplemented with the wash solution and incubated for 30 min. For agonist assays the samples were injected onto the cells pre-incubated with the wash solution. Changes in cellular fluorescence were monitored using a NOVOStar fluorescence plate reader (BMG Labtech). Data were normalized to the maximum change in fluorescence induced by TRPM8 agonist icilin (50 µM). AMTB hydrochloride (20 µM), a TRPM8 channel blocker, was used as a positive control in the antagonist assay. The IC_50_ value for the sample was calculated using nonlinear regression in GraphPad 7 (GraphPad Software).

### Two-electrode voltage-clamp experiments

cRNA of hK_V_3.1, hK_V_7.2, and hK_Ca_1.1 were prepared from linearized plasmids using the mMESSAGE mMACHINE T7 kit (Ambion) according to manufacturer’s instructions. RNA quality and concentrations were assessed by UV spectroscopy (NanoDrop, Thermo Scientific) and gel electrophoresis. The prepared cRNA for K_V_7.2 (10 ng), K_Ca_1.1 (5 ng) or K_V_3.1 (0.5 ng) was injected into *Xenopus laevis* oocytes (EcoCyte Bioscience).

Currents were recorded using a Dagan CA-1B amplifier after 1-3 days of incubation (19 °C). Data acquisition was performed with the Pulse software (HEKA Elektronik). Borosilicate glass electrodes (Module Ohm) were pulled on a DMZ-Universal Puller (Zeitz Instruments) and filled with 2 M KCl. Oocytes were superfused with Kulori solution (NaCl (90 mM), KCl (4 mM), MgCl_2_ (1 mM), CaCl_2_ (1 mM), HEPES (5 mM), pH=7.4 at r.t.).

The experiments comprised a baseline and a compound (DIMTA) perfusion period. During the baseline period, the membrane potential was depolarized every 10th s by an online voltage protocol. After 10 depolarizations, the oocyte was subjected to a voltage step protocol in order to generate a current-voltage (IV) relationship. Immediately after this, the solution was changed to Kulori solution containing DIMTA (5 µM). During the incubation with DIMTA, the membrane potential was depolarized every 10th second by an online voltage protocol. After the 30^th^ depolarization, the oocyte was subjected to a voltage-step protocol in order to generate an IV-relationship in the presence of DIMTA. We generated the current-voltage relationship (I/V-curve) by plotting the mean of the last 20 ms of each voltage step.

### Patch clamp experiments

To investigate the effects of DIMTA on K_V_3.1 (isoform b), 0.6 µg K_V_3.1b plasmid and 0.1 µg eGPF were transfected into HEK-293 cells using Lipofectamine 2000 (Invitrogen Corporation), and recordings were performed after 3 days of incubation at 37 °C. The effects of DIMTA on K_Ca_1.1 and K_V_11.1 were studied using HEK293 cells and Chinese Hamster Ovary (CHO) constitutively expressing human K_Ca_1.1 and K_V_11.1, respectively.

For patch-clamp experiments, the cells were washed with PBS, detached from the culture flask using trypsin (ThermoFisher Scientific) or Detachin™ (Amsbio) and seeded onto glass coverslips for manual patch clamping or automatically loaded onto disposable single-hole Qplates (Biolin Scientific) for automated patch clamping. For K_Ca_1.1 inside-out experiments, the coverslips were coated with 50 mg/L poly-L-lysine at 37 °C overnight in order to get better attachment of the cells to the coverslip. The coverslips were placed into a custom-made perfusion recording chamber with ∼1 ml/min continuous superfusion.

The recordings were performed at room temperature. Only cells that maintained a high membrane resistance seal above 1 GΩ and had a maximum serial resistance (Rs) of 10 MΩ were used for subsequent analysis. The serial resistance was compensated for 50 to 80% during the experiment. Data was acquired using a Multiclamp 700B amplifier (Axon Instruments). The analogue output signals were digitized and recorded after low pass filtering at 2.9 kHz through Digidata 1322A/1440A pClamp 10.2 (Molecular Devices). The patch-clamp borosilicate glass micropipettes were pulled by a horizontal DMZ universal puller (Zeitz Instruments). The electrodes had resistances between 1.5 and 3 MΩ when filled with solution.

The extracellular solution for all experiments contained: NaCl (145 mM), CaCl_2_ (2 mM) MgCl_2_ (1 mM), KCl (4 mM), glucose (10 mM), and HEPES (10 mM), pH 7.4 adjusted with NaOH. The intracellular solution contained: CaCl_2_ (5.17 mM), MgCl_2_ (1.42 mM), KOH/EGTA (31.25/10 mM), KCl (114 mM), KOH (9 mM) and HEPES (10 mM), pH adjusted with HCl to 7.4, resulting in a free calcium concentration of 100 nM.

### K_Ca_1.1 currents

K_Ca_1.1 currents were activated by ramping the membrane potential from −80 mV to +50 mV (200 ms) from a holding potential of −80 mV. The protocol was repeated every 2 s. The cell was stabilized for 1 to 5 min and then subjected to the following perfusion protocol: 1 min recording in extracellular solution (baseline); and 2 min in 1, 3, 10, 30, 100, 300 µM DIMTA. In some experiments we switched back to the extracellular solution to observe if the effect of DIMTA could be washed off.

### K_V_3.1 currents

To study the effect of DIMTA on K_V_3.1 four different voltage protocols were applied. (1) IV protocol: the cells were held at −80 mV for 4800 ms followed by a 200 ms depolarizing pulses from −80 mV to +60 mV in +10 mV increments. (2) Online protocol: every 4^th^ s the cells were activated to +10 mV for 100 ms from a holding potential of −80mV. (3) Inactivation protocol: the cells were held at membrane potentials between −100 to −10 mV for 30 s, followed by a depolarization to +40 mV. For each trial, the cell was held at −80 mV for 1 min to allow the ion channel to recovery from inactivation. (4) Instantaneous IV curve: to generate plots of the voltage dependence of activation (g/gmax) the reversal potential (Vrev) was established by depolarizing the cell membrane to 40 mV for 200 ms, followed by a voltage steps to −60 to −120 for 5 s in 5 mV increments.

After stabilization of the K_V_3.1 current the cells were subjected to the following perfusion protocol: 1 min recording in extracellular solution (baseline), and subsequently 2 min in 1, 3, 10, 30, 100 µM DIMTA. During these perfusion periods the online voltage protocol was continuously applied. At the end of each perfusion period we ran the IV voltage protocol. Finally, in some cells, to investigate recovery from inhibition we switched back to extracellular solution.

In a separate set of experiments the effect of DIMTA on steady state inactivation was investigated. Following stabilization, the cells were subjected to the inactivation voltage protocol during normal extra cellular solution. We switched the solution to DIMTA, and once the effect of the compound had stabilized, we repeated the inactivation voltage protocol in the presence of the compound.

### Kv11.1 currents

We used the automatic patch clamp system QPatch-16 (Sophion) and disposable single-hole Qplates (Sophion) to investigate the effect of DIMTA on K_V_11.1 channels. The system allows for automatic giga seal formation, whole-cell formation, access resistance compensation, liquid application, and recording. CHO cells constitutively expressing K_V_11.1 channels were held at –90 mV and currents activated by stepping to +20 mV from −80 mV for 2 s and then to –50 mV for 2 s in order to record the tail current. The time between each depolarization was 7 s. Data were sampled at 10 kHz, four-order Bessel filter, cut-off frequency 3 kHz and 70 % Rs compensation. The perfusion protocol used was: baseline recordings in extracellular solution followed by application of increasing concentration of DIMTA (3, 10, 30, 60, 100, 300 µM).

### *In vivo* cold plate assay

Behavioral testing was conducted according to procedure described previously.^21, 33^ In brief, male adult (6-8 weeks old) CD1 mice were injected i.p. with DIMTA (1 mg/kg, 10 mg/kg), or vehicle (1% DMSO in saline) 30 min prior to behavioral testing. Mice were assessed in a cold-plate test chamber (IITC, Inc. Life Science) in which the temperature was decreased from room temperature at a rate of 10 °C per minute. The testing was stopped when pain-related behaviors including lifting and licking or shaking of the fore-paws was observed. Alternating lifting of a single forepaw was not scored as a response. The latency and temperature of the first sign of pain response were recorded. Thirteen mice were used per treatment group and behavioral test measurement was done once per mouse. Data were evaluated with one-way Anova and Dunnett’s multiple comparisons test using Graphpad Prism (Graphpad Software).

### Data analysis

For two-electrode voltage clamp, data were analyzed using the PatchMaster software (HEKA), and GraphPad Prism (GraphPad Software). Patch clamp data and automated patch clamp data were analyzed using Clampfit 10.7 software, QPatch software and GraphPad Prism. All values are expressed as means ± SEM. All electrophysiological recordings were analyzed using one-way or two-way analysis of variance (ANOVA) followed by Tukey’s or Sidak multiple comparison test. *p* < 0.05 was considered significant.

IV and IV-peak curves were constructed from the IV protocol, and all data were plotted against the corresponding membrane potentials. The IV and IV-peak curves were constructed from measurements at the end of the depolarization and at the peak current amplitude, respectively.

Voltage dependence of activation was studied using normalized conductance (g/gmax) curves. These curves were generated from the I-V curves and the reversal potential (Vrev). The latter was experimentally found using the instantaneous IV voltage protocol. We calculated the conductance for each test potential using the formula: G = I/(Vm-Vrev), where I is current amplitude and Vm is membrane potential. The resulting data were then fitted with a Boltzmann function: g/gmax = 1/(1 + exp[− (V − V1/2)/k]), where Gmax is the maximal conductance, V is the membrane potential, V1/2 is the potential at which the value of the relative conductance is 0.5, and k is the slope factor.

To determine the voltage dependence of inactivation the peak current at +40 mV was normalized to the maximal K_V_3.1 current and plotted as a function of the preceding holding potential. The V_50_ of inactivation was determined by fitting a Boltzmann sigmoidal function to the individual inactivation curves.

The concentration-response curves were constructed by the plotting the log(concentration) against the current density. Individual IC_50_ values for each experiment were calculated using the equation:

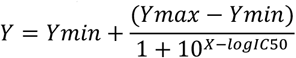

where X is the log of concentration of 12114 and Y is the measured current. The bottom a of the curve was constrained to 0.

### Estimation of indirect effects (IDE) in calcium imaging experiments with DRG neurons

IDE of the compound was estimated using repeated 15 s pulses of K^+^ (25 mM). Each pulse elicits a calcium response from all neurons in the experiment. The magnitude of the response was measured as the maximum area under the curve in any 15 s interval during incubation with K^+^ (*Kauc*). This maximal response is highly reproducible for each cell. The deviation of this response after a 5-min incubation with each compound was used to estimate the indirect effect of the compound. A linear model (lm function in R) was applied to each neuron: lm(*Kauc* ∼ *linear* + c1 + c2 + c3 + c4 + c5)

Where *Kauc* is the statistic described above for each of the 16 K^+^ pulses, *linear* is the sequential trend coded as sequential integers, c1, c2, c3, c4 and c5 are indicator variables for the K^+^ pulse that is under the influence of the specific compounds (DIMTA (2.5 µM), DIMTA (25 µM), TEA (10 mM), nicardipine (400 nM), TEA (10 mM) + nicardipine (400 nM). The Tstat values from the coefficients matrix were taken as estimates of the magnitude and direction of IDE.

### Estimation of direct effects (DE) in calcium imaging experiments with DRG neurons

DE of the compound was estimated by comparing the compound incubation interval to control interval incubations with only DRG observation buffer. The same statistics were calculated as detailed above. In this case the Tstat values from the linear regression are interpreted as the DE of the compound.

### Estimating significance for multiple tests

Each experiment gave results for 500 – 1000 neurons. To control for false positives and set appropriate thresholds for significance we used Monte Carlo simulations. The Tstats were estimated for all cells, recorded, and considered as the null distribution. The Tstat estimates from 100 Monte Carlo simulations were used to establish the thresholds for single test cell significance as well as whole experiment significance. The average 95% interval from the 100 Monte Carlo simulations was used as the threshold for an individual cell. Cells that exceeded the lower bound were considered blocked. Those that exceeded the higher bound were considered amplified. The per-experiment significance was estimated as the fraction of Monte Carlo simulations that were different from the actual data at a ks.test threshold of 0.01. The nature of the IDE is evident when comparing the actual Tstat distribution to the null distributions as empirical cumulative distribution function (ecdf) curves (**Figure S1**).

### Associations Between Effects

Correlations between effects using the Tstats were estimated using the cor.test function of R.

### Estimation of association between gene expression counts and cell TRPM8 class

Using single cell RNA seq data from DRG neurons,^17^ a logistic regression analysis was used to estimate the association between gene expression counts and cell TRPM8 class membership.

## AUTHOR INFORMATION

### Author Contributions

The manuscript was written through contributions of all authors. All authors have given approval to the final version of the manuscript.

### Notes

The authors declare no competing financial interest.

## ACKNOWLEDGMENT

Assistance from the Ministry of Environment, Climate Change, Disaster Management, and Meteorology (Solomon Islands) and Solomon Islands National University is gratefully acknowledged.

The graphical abstract was created with biorender.com.

