## Supporting Information for "The tunicate metabolite 2-(3,5-diiodo-4-methoxyphenyl)ethan-1-amine targets ion channels of vertebrate sensory neurons"

### SUPPORTING FIGURES

**Figure S1.** Census of effects elicited by DIMTA (2.5 μM) on different subsets of DRG neurons.
**Figure S2.** Constellation pharmacology shows the effects of DIMTA on calcium and sodium channels in low-threshold cold thermosensors (LTCTs).
**Figure S3.** Inside-out recordings on $hK_{Ca}1.1$ expressed in HEK-293 cells.
**Figure S4.** Whole-cell electrophysiology with $hK_{Ca}1.1$.
**Figure S5.** Effects of increasing concentrations of DIMTA on $K_V11.1$ expressed in HEK-293 cells.
**Figure S6.** Evaluation of DIMTA as an inhibitor of human transient receptor potential melastatin 8 (TRPM8) overexpressed in HEK-293 cells.
**Figure S7.** Spectroscopic, spectrophotometric, and chromatographic analysis of DIMTA.

### SUPPORTING TABLES

**Table S1.** K channel genes (KCN) associated with TRPM8 expressing cells.
**Table S2**. Raw data for in vivo cold plate assay.

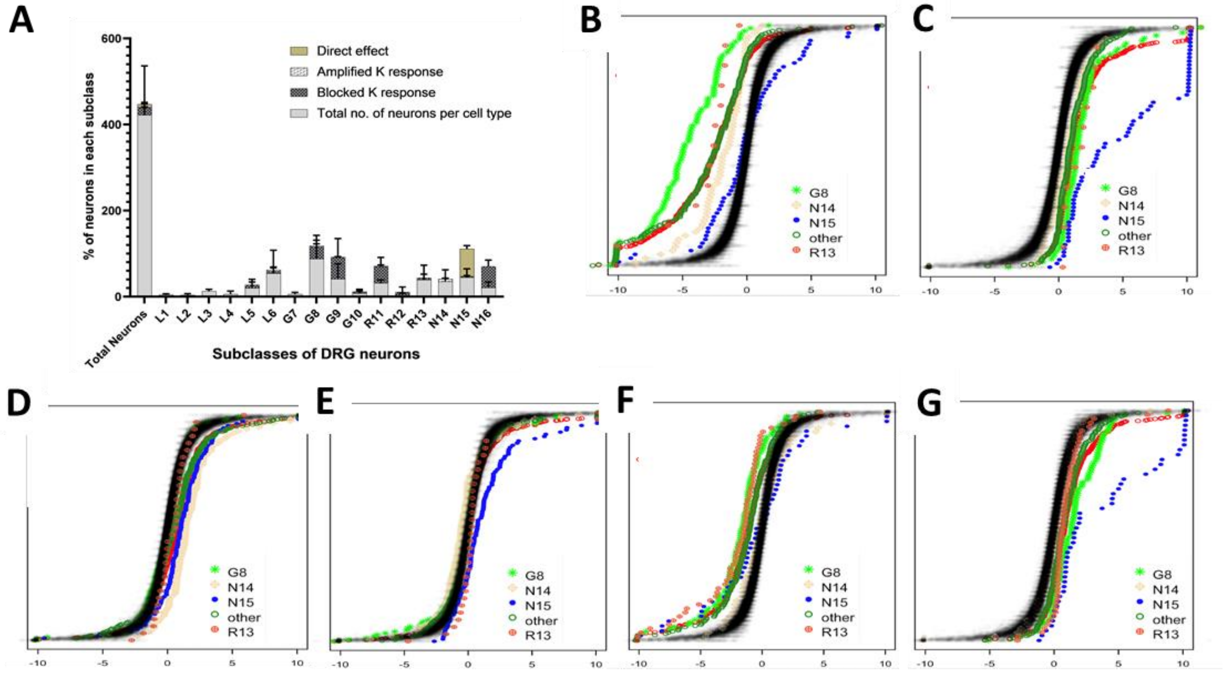


**Figure S1.** Census of effects elicited by DIMTA (2.5 µM) on different subsets of DRG neurons. **(A)** Bar plot summarizing the effects of DIMTA in calcium imaging experiments (n = 3). Block/amplification of the $Ca^{2+}$ signal, or direct effects to the $Ca^{2+}$ baseline with sample incubation were scored. The effects for each subtype (x-axis) are represented as the mean % of neurons in each subclass ± standard deviation (y-axis). **(B-G)** Statistical analysis to estimate indirect (IDE, panels **B**, **D** and **F**) and direct effects (DE, panels **C**, **E**, **G**) for each calcium imaging experiment. The nature of the IDE and DE is compared to the actual Tstat distribution to the null distribution as empirical cumulative distribution function (ecdf) curves. The x-axis, the value of -10 to 10 indicates the IDEs block to amplification in **(B)**, **(D)**, and **(F)**. For **(C)**, **(E)**, and **(G)** x-axis value > 0 indicates DE**.** The x-axis value = 0 indicates no effect. The y-axis indicates the relative number of cells that fall below the given value (ecdf), from 0 (bottom) to 1.0 (top). **(B)** IDE and **(C)** DE estimation in experiment 1. **(D)** IDE and **(E)** DE estimation in experiment 2. **(F)** IDE and **(G)** DE estimation in experiment 3. N15 represents the thermosensors while G8, N14, R13, and other are neurons belonging to different subclasses.

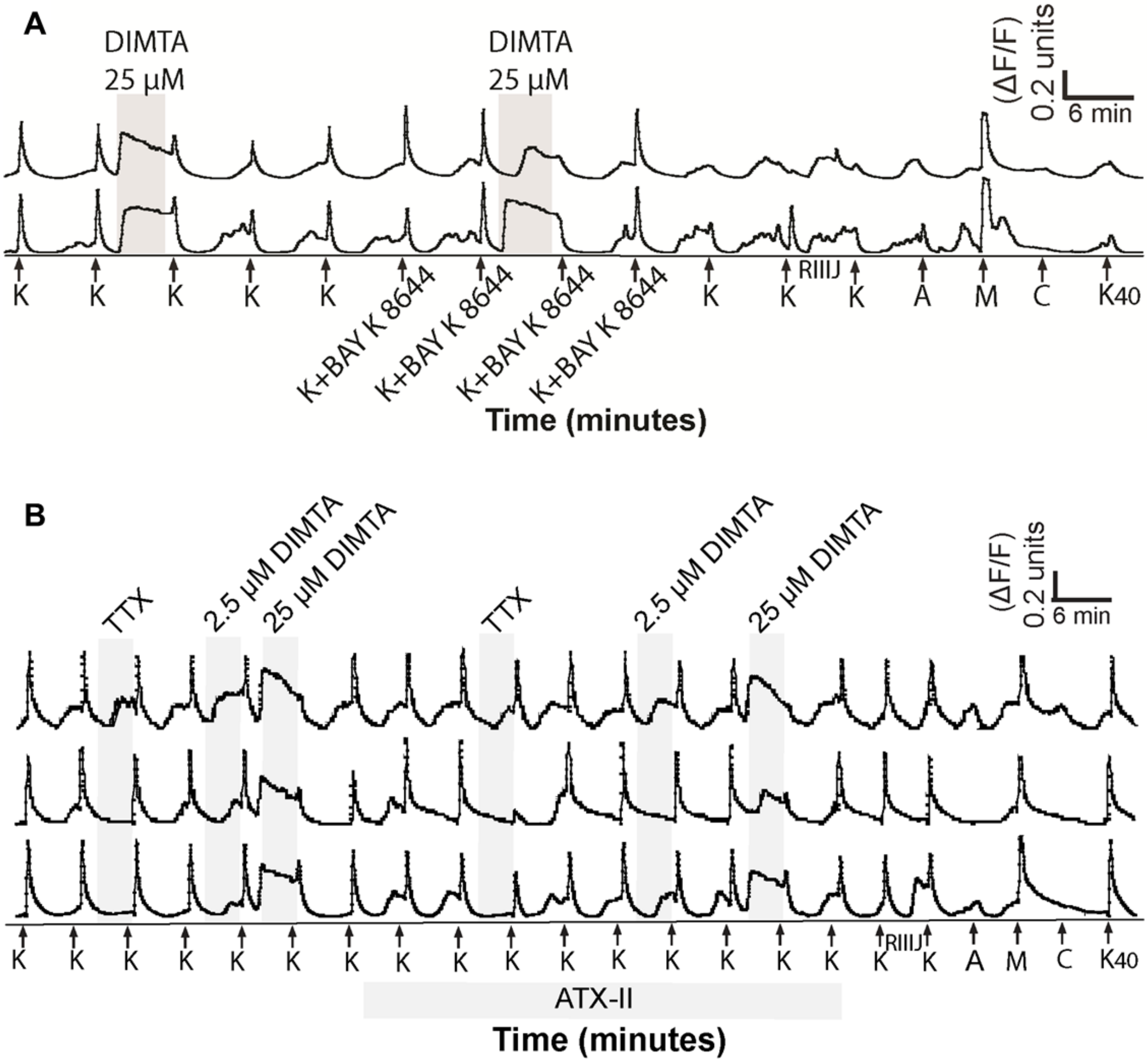


**Figure S2.** Constellation pharmacology shows the effects of DIMTA on calcium and sodium channels in low-threshold cold thermosensors (LTCTs). **(A)** and **(B)** represent constellation pharmacology using DIMTA. Each trace follows the Fura2-AM fluorescence min/max normalized ratio at 340/380 nm (a measure of relative intracellular calcium levels) (*y*-axis) of a single cell out of 500-1000 neurons in a single experiment. Chemicals were added over the experiment time (*x*-axis), and the cellular responses were recorded. Arrows indicate 15-s application of potassium chloride (K, 20 mM) or co-application of K and Bay K 8644 (200 nM). The highlighted region indicates the period of DIMTA or pharmacological agent incubation. Sequential application of a set of pharmacological ligands was used to identify neuronal cell populations: K, RIIIJ (1 µM), allyl isothiocyanate (A, 100 µM), menthol (M, 400 µM), and capsaicin (C, 300 nM). Potassium chloride 40 mM ($K_{40}$) was applied at the end of the experiment to determine the viability of the neurons. **(A)** Selected traces from LTCTs illustrating the effects of DIMTA (25 µM) on calcium ($Ca^{2+}$) channels. Bay K 8644 was co-applied with K to activate L-type $Ca^{2+}$ channels. **(B)** Selected traces from LTCTs illustrating the effects of DIMTA (2.5 µM and 25 µM) and tetrodotoxin (TTX, 1 µM), a VGSC blocker on sodium ($Na^+$) channels. *Anemonia viridis* toxin 2 (ATX-II, 100 nM) was applied to delay $Na^+$ channel inactivation.

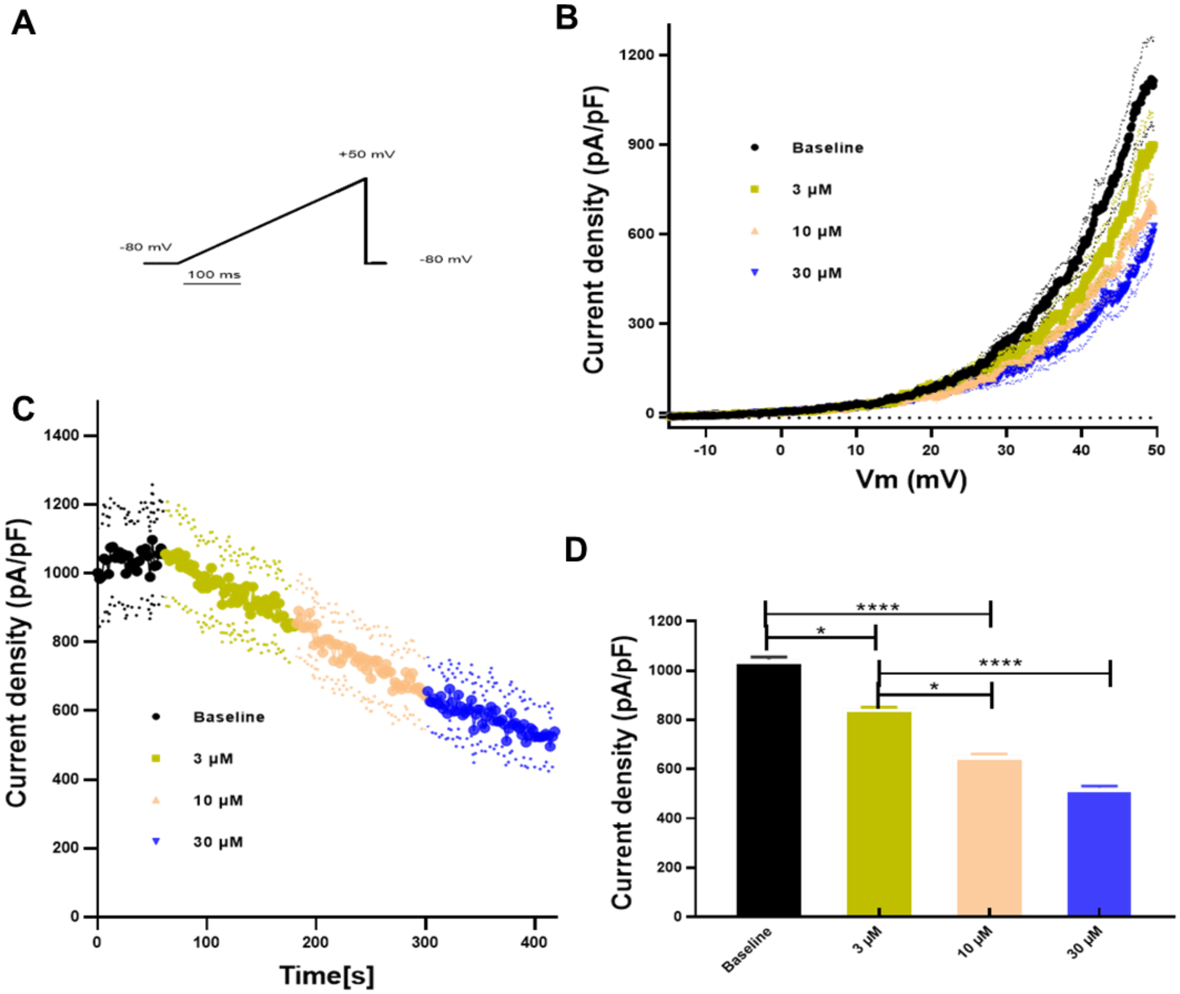


**Figure S3.** Inside-out recordings on $hK_{Ca}1.1$ expressed in HEK-293 cells. **(A)** Currents were elicited every 2 s using the voltage-clamp protocol shown. **(B)** Mean current-voltage relationships and **(C)** current-time plots before and after application of increasing concentrations (3 μM, 10 μM, 30 μM) of DIMTA. **(D)** Bar graph summarizing the effect of increasing concentrations of DIMTA on $K_{Ca}1.1$ currents measured at +50 mV. Data are presented as mean ± SEM, n=10.

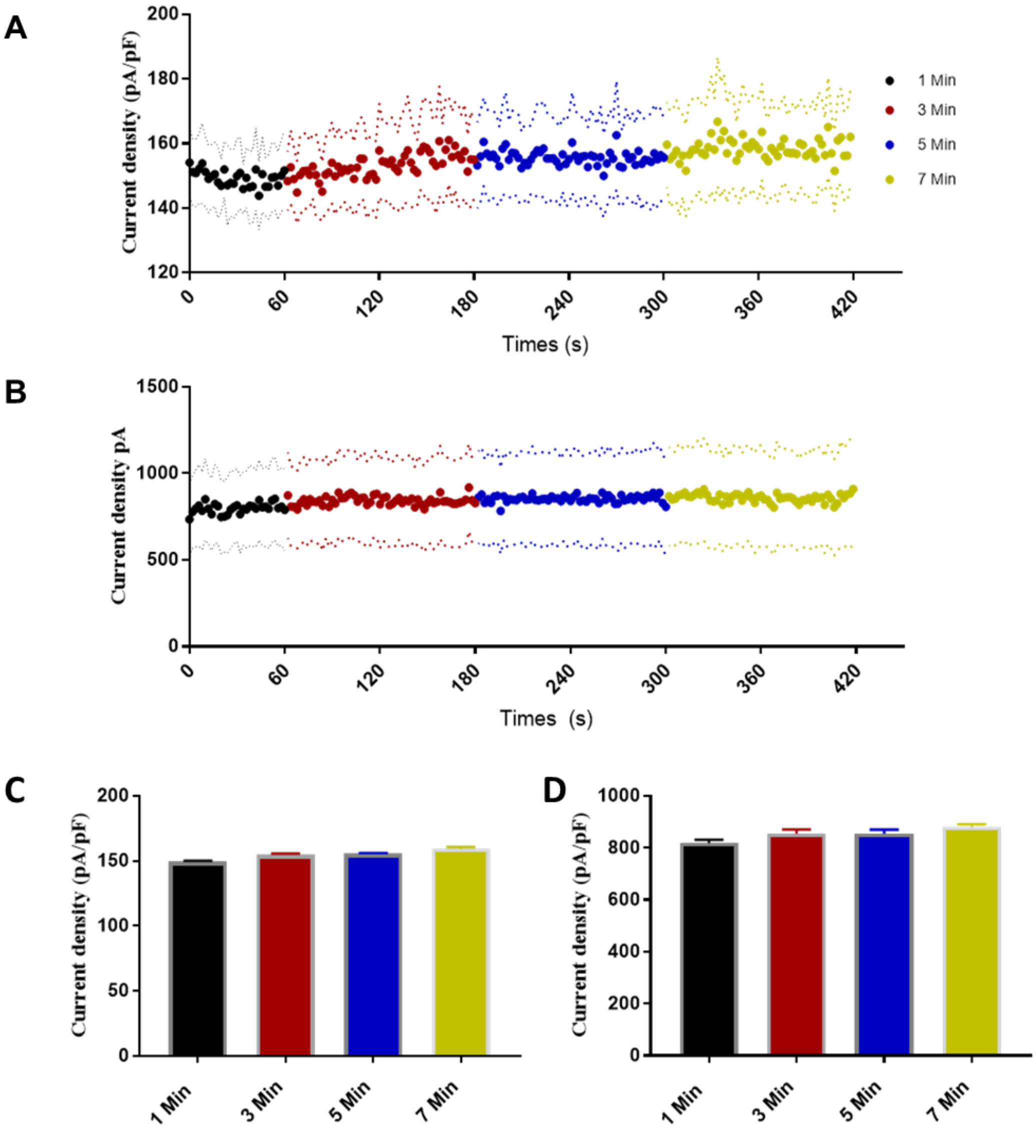


**Figure S4.** Whole-cell electrophysiology with h$K_{Ca}$1.1. **(A)** Time matched control experiment on h$K_{Ca}$1.1 with whole-cell configuration, n=5. **(B)** Time dependent control experiment of inside-out recordings of $K_{Ca}$1.1, n=11. **(C)** Bar graph summarizing the control experiment using $K_{Ca}$1.1 with whole-cell configuration, currents measured at +50 mV, bars represent mean ± SEM, n= 5. **(D)** Bar graph summarizing the control experiment using $K_{Ca}$1.1 with inside-out configuration, currents measured at +50 mV, bars represent mean ± SEM, n= 11.

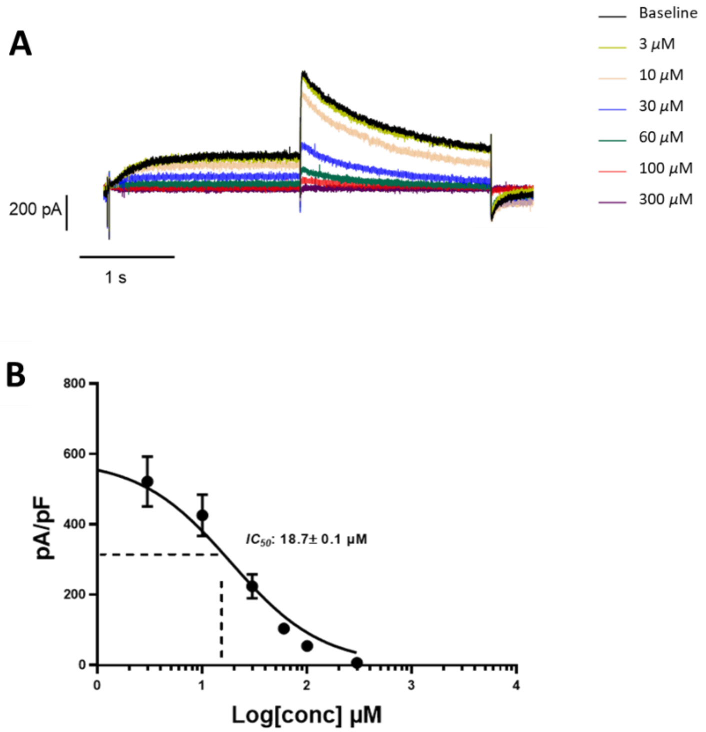


**Figure S5.** Effects of increasing concentrations of DIMTA on $K_V11.1$ expressed in HEK-293 cells. **(A)** Representative current recording of $K_V11.1$ before and after application of increasing concentrations of DIMTA. **(B)** Concentration response relationship for DIMTA inhibition of $K_V11.1$ channel currents. Data are presented as mean ± SEM, n=5-7.

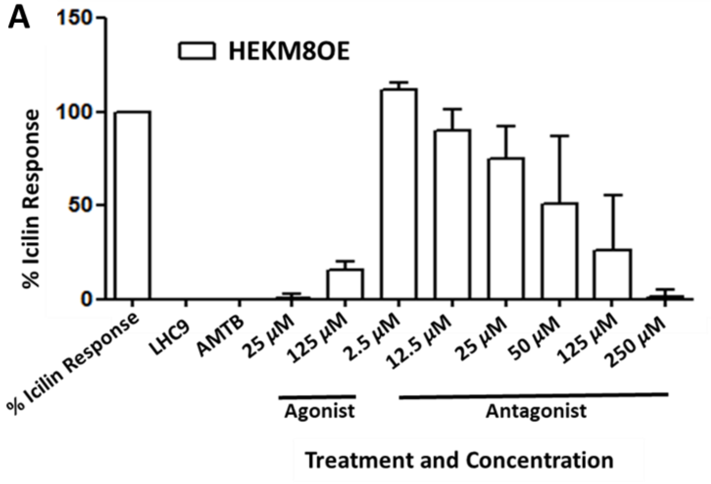


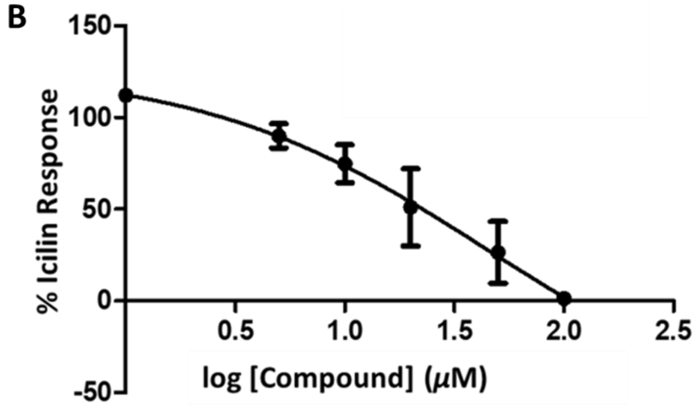


**Figure S6.** Evaluation of DIMTA as an inhibitor of human transient receptor potential melastatin 8 (TRPM8) overexpressed in HEK-293 cells. **(A-B)** Calcium flux in HEK-293 cells was induced by addition of icilin, a synthetic agonist of TRPM8. Data are normalized to the maximum fluorescence change elicited by the positive control (50 µM icilin). **(A)** Agonist and antagonist activity (x-axis) of DIMTA is normalized to icilin response and presented as % Icilin response (y-axis). LHC9 is the media/buffer used in the calcium imaging assays. AMTB (20 µM) is a commercially available TRPM8 antagonist. **(B)** The concentration response inhibition of hTRPM8 by DIMTA represented as % Icillin Response (y-axis) vs log of DIMTA concentration (x-axis). Values are the mean ± SEM from triplicate wells. The $IC_{50}$ is 112 µM with a Hill slope of -0.66.

A

B

C

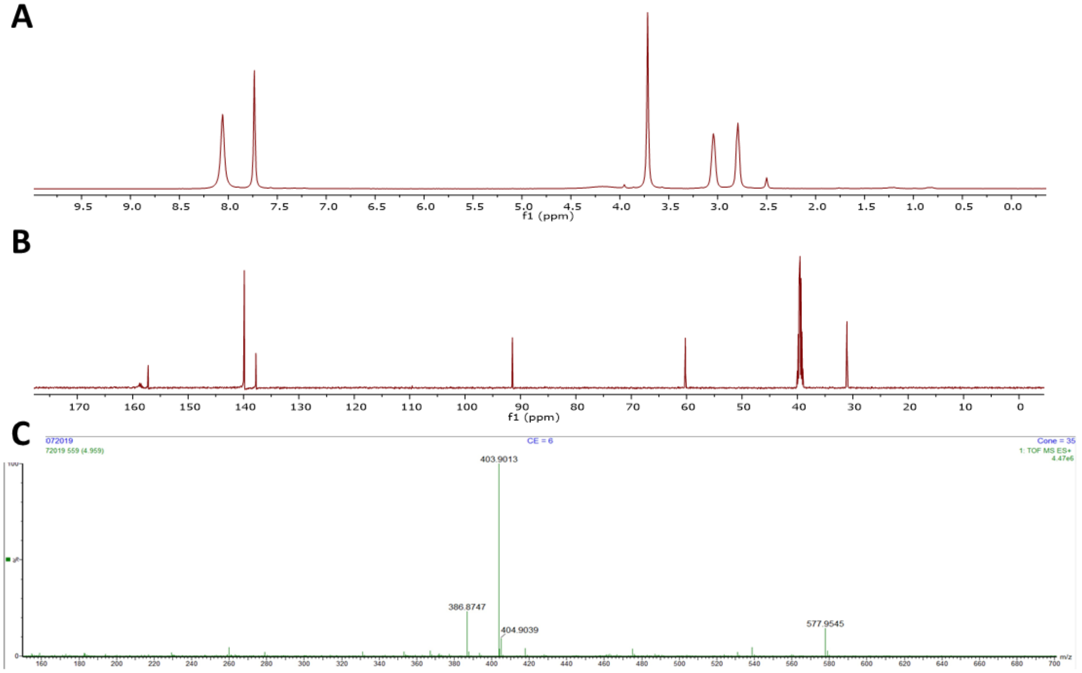


D

**Single Mass Analysis**
Tolerance = 5.0 PPM / DBE: min = -1.5, max = 50.0
Element prediction: Off
Number of isotope peaks used for i-FIT = 3

Monoisotopic Mass, Even Electron Ions
3396 formula(e) evaluated with 4 results within limits (all results (up to 1000) for each mass)
Elements Used:
C: 0-500 H: 0-1000 N: 0-200 O: 0-200 23Na: 0-1 127I: 0-2

1-211-4_08072019 CE = 6 Cone = 35
1-211-4_08072019 559 (4.959) 1: TOF MS ES+
4.47e+006

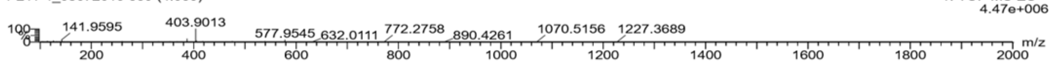


| Mass | Calc. Mass | mDa | PPM | DBE | i-FIT | Norm | Conf(%) | Formula |
|---|---|---|---|---|---|---|---|---|
| Minimum: | | | | -1.5 | | | | |
| Maximum: | | 5.0 | 5.0 | 50.0 | | | | |
| 403.9013 | 403.9008 | 0.5 | 1.2 | 3.5 | 709.7 | 0.002 | 99.85 | C9 H12 N O 127I2 |
| | 403.9016 | -0.3 | -0.7 | 10.5 | 716.8 | 7.034 | 0.09 | C10 H3 N3 O7 127I |
| | 403.9005 | 0.8 | 2.0 | 12.5 | 717.1 | 7.374 | 0.06 | C9 N7 O3 23Na 127I |
| | 403.9032 | -1.9 | -4.7 | 11.5 | 720.1 | 10.421 | 0.00 | C13 H4 N O5 23Na 127I |

E

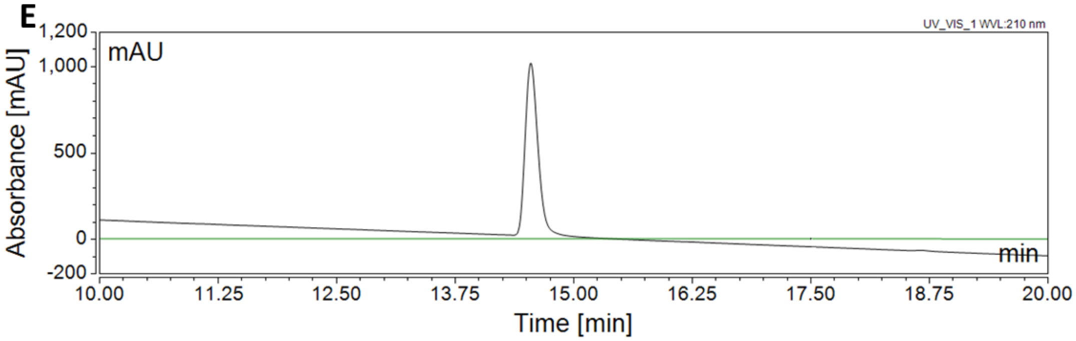


**Figure S7.** Spectroscopic, spectrophotometric, and chromatographic analysis of DIMTA. **(A)** $^{1}H$ NMR (500 MHz) and **(B)**$^{13}C$ NMR (125 MHz spectra of DIMTA in DMSO-$d_6$. **(C)-(D)** (+)-HRESITOFMS of DIMTA showing m/z 403.9013 $[M+H]^+$ (calculated for $C_9H_{12}NOI_2$, 403.9003). **(E)** Chromatographic profile of DIMTA using an analytical Eclipse Plus $C_{18}$ column (3.5 μm; 4.6 x 150 mm) and eluted with a linear gradient from 5% to 100% $CH_3CN$ in $H_2O$ (0.1% TFA) over 24 min at 0.5 mL/min flow rate. The UV absorbance was monitored at 210 nm.

**Table S1.** K channel genes (*KCN*) associated with TRPM8 expressing cells.

| Gene | Mean | glt.z | frac.0M8 | frac.1M8 | sLog10p |
|---|---|---|---|---|---|
| *KCNK2* | 0.992 | 18.7 | 0.290 | 0.750 | 92 |
| *KCNN3* | 0.022 | 22.5 | 0.009 | 0.213 | 89 |
| *KCNMB2* | 0.302 | 19.6 | 0.154 | 0.539 | 80 |
| *KCNK3* | 0.543 | 12.6 | 0.212 | 0.467 | 34 |
| *KCNN2* | 0.145 | 7.4 | 0.109 | 0.221 | 11 |
| ***KCNMA1*** | **0.631** | **6.9** | **0.378** | **0.535** | **11** |
| *KCNQ1OT1* | 0.641 | 6.8 | 0.401 | 0.557 | 11 |
| *KCNA2* | 1.751 | 6.4 | 0.611 | 0.756 | 10 |
| *KCNH8* | 0.087 | 6.9 | 0.069 | 0.154 | 10 |
| *KCNC2* | 0.696 | 6.2 | 0.389 | 0.531 | 9 |
| *KCNA4* | 0.536 | 4.3 | 0.352 | 0.447 | 5 |
| *KCNC4* | 2.397 | 4.1 | 0.568 | 0.662 | 4 |
| ***KCNC1*** | **0.385** | **4.1** | **0.221** | **0.301** | **4** |
| *KCND2* | 0.023 | 4.4 | 0.019 | 0.049 | 4 |
| *KCNH7* | 1.252 | 2.6 | 0.519 | 0.580 | 2 |
| *KCNAB2* | 7.182 | 0.1 | 0.989 | 1.000 | 2 |
| *KCNJ4* | 0.053 | 2.4 | 0.043 | 0.066 | 2 |
| *KCNB2* | 5.636 | 1.9 | 0.958 | 0.975 | 1 |
| ***KCNQ2*** | **0.568** | **1.6** | **0.349** | **0.383** | **1** |
| *KCNC3* | 0.299 | 0.1 | 0.218 | 0.219 | 0 |
| *KCNJ10* | 0.097 | -0.1 | 0.077 | 0.076 | 0 |
| *KCNH6* | 0.060 | -1.0 | 0.058 | 0.047 | 0 |
| *KCNAB3* | 0.014 | -1.3 | 0.013 | 0.006 | -1 |
| *KCNMB4OS* | 0.112 | -1.4 | 0.099 | 0.080 | -1 |
| *KCNJ16* | 0.012 | -1.5 | 0.012 | 0.004 | -1 |
| *KCNJ12* | 0.471 | -1.4 | 0.290 | 0.260 | -1 |
| *KCNF1* | 0.044 | -1.5 | 0.041 | 0.027 | -1 |
| *KCNH2* | 0.240 | -1.5 | 0.164 | 0.139 | -1 |
| *KCNT1* | 1.265 | -16.1 | 0.582 | 0.061 | -127 |
| *KCNIP3* | 1.343 | -17.9 | 0.629 | 0.113 | -119 |
| *KCNAB1* | 1.438 | -12.7 | 0.470 | 0.031 | -103 |
| *KCNQ2* | 0.511 | -0.1 | 0.348 | 0.000 | -88 |
| *KCNA6* | 1.207 | -16.7 | 0.633 | 0.203 | -79 |
| *KCNMB1* | 1.056 | -10.5 | 0.376 | 0.020 | -79 |
| *KCNIP2* | 1.285 | -13.3 | 0.448 | 0.059 | -79 |
| *KCNH1* | 0.533 | -11.8 | 0.379 | 0.055 | -60 |
| *KCND1* | 1.574 | -15.6 | 0.698 | 0.332 | -59 |
| *KCND3* | 1.632 | -13.0 | 0.517 | 0.189 | -48 |
| *KCNN1* | 0.413 | -10.2 | 0.308 | 0.041 | -48 |
| *KCNK12* | 0.404 | -9.6 | 0.279 | 0.037 | -42 |
| *KCNQ5* | 0.301 | -6.8 | 0.199 | 0.008 | -39 |
| *KCNV1* | 0.244 | -4.5 | 0.152 | 0.002 | -32 |
| *KCNJ3* | 0.316 | -8.4 | 0.227 | 0.039 | -30 |
| *KCNS1* | 0.211 | -6.3 | 0.153 | 0.010 | -27 |
| *KCNT2* | 0.155 | -0.1 | 0.108 | 0.000 | -23 |
| *KCNK13* | 0.241 | -7.6 | 0.198 | 0.043 | -22 |
| *KCNA1* | 0.524 | -7.7 | 0.263 | 0.098 | -18 |
| *KCNK1* | 0.270 | -6.2 | 0.171 | 0.057 | -13 |
| *KCNIP1* | 0.146 | -5.7 | 0.120 | 0.027 | -13 |
| *KCNK10* | 0.058 | -3.3 | 0.055 | 0.002 | -10 |
| *KCNK18* | 0.084 | -4.6 | 0.075 | 0.010 | -10 |
| *KCNQ3* | 0.253 | -4.9 | 0.101 | 0.029 | -9 |
| *KCNS3* | 0.541 | -5.7 | 0.335 | 0.209 | -9 |
| *KCNG4* | 0.040 | -0.1 | 0.034 | 0.000 | -7 |
| *KCND3OS* | 0.058 | -3.8 | 0.057 | 0.012 | -6 |
| *KCNJ2* | 0.038 | -2.9 | 0.036 | 0.002 | -6 |

mean=mean expression across 11,139 cells. glt.z = zscore from logistic regression onto binary TrpM8. frac.0M8 = fraction of cells expressing the gene (> 0) in TrpM8 negative, frac.1M8 = fraction of cells expressing the gene (>0) in TrpM8 positive cells. sLog10p is signed log10(fisher.test$pvalue).

**Table S2**. Raw data for *in vivo* cold plate assay.

| Vehicle* | | 1 mg/kg DIMTA | | 10 mg/kg DIMTA | |
|---|---|---|---|---|---|
| Time (sec) | Temperature °C | Time (sec) | Temperature °C | Time (sec) | Temperature °C |
| 86.72 | 9.5 | 162.95 | 0.2 | 67.65 | 12.7 |
| 72.07 | 12 | 101.93 | 7 | 156.59 | 0.2 |
| 57.68 | 14.3 | 96.37 | 7.9 | 75.24 | 11.4 |
| 78.35 | 10.9 | 89.97 | 9 | 139.6 | 1.4 |
| 70.69 | 12.1 | 87.26 | 9.4 | 92.11 | 8.7 |
| 104.37 | 6.7 | 141.3 | 1.1 | 144.38 | 0.5 |
| 141.06 | 1.2 | 106.49 | 6.4 | 93.53 | 8.5 |
| 109.01 | 6 | 87.91 | 8.3 | 168.88 | 0.3 |
| 99.52 | 7.3 | 121.36 | 4.1 | 140.13 | 1.4 |
| 105.06 | 6.6 | 141.79 | 1 | 107.94 | 6.1 |
| 106.01 | 6.1 | 156.81 | 0.3 | 132.48 | 2.4 |
| 111.22 | 5.6 | 89.94 | 9 | 108.89 | 6 |
| 105.25 | 6.5 | 109.08 | 6 | 159.83 | 0.3 |

*1% DMSO in normal saline solution
